# Remodeling of yeast vacuole membrane lipidomes from the log (1-phase) to stationary stage (2-phases)

**DOI:** 10.1101/2022.10.11.511736

**Authors:** John Reinhard, Chantelle L. Leveille, Caitlin E. Cornell, Alexey J. Merz, Christian Klose, Robert Ernst, Sarah L. Keller

**Affiliations:** Medical Biochemistry and Molecular Biology, Medical Faculty, Saarland University, Homburg, Germany; PZMS, Center for Molecular Signaling, Medical Faculty, Saarland University, Homburg, Germany; Department of Chemistry, University of Washington, Seattle, WA, USA; Department of Biochemistry, University of Washington, Seattle, WA, USA; Lipotype GmbH, Am Tatzberg 47, Dresden, Germany

**Author notes:** These authors contributed equally to this work (John Reinhard, Chantelle L. Leveille).

**Keywords:** yeast, vacuole, membrane, lipidome, phase separation

## Abstract

Upon nutrient limitation, budding yeast of *Saccharomyces cerevisiae* shift from fast growth (the log stage) to quiescence (the stationary stage). This shift is accompanied by liquid-liquid phase separation in the membrane of the vacuole, an endosomal organelle. Recent work indicates that the resulting micron-scale domains in vacuole membranes enable yeast to survive periods of stress. An outstanding question is which molecular changes might cause this membrane phase separation. Here, we conduct lipidomics of vacuole membranes in both the log and stationary stages. Isolation of pure vacuole membranes is challenging in the stationary stage, when lipid droplets are in close contact with vacuoles. Immuno-isolation has previously been shown to successfully purify log-stage vacuole membranes with high organelle specificity, but it was not previously possible to immuno-isolate stationary stage vacuole membranes. Here, we develop Mam3 as a bait protein for vacuole immuno-isolation, and demonstrate low contamination by non-vacuolar membranes. We find that stationary stage vacuole membranes contain surprisingly high fractions of phosphatidylcholine lipids (∼50%), roughly twice as much as log-stage membranes. Moreover, in the stationary stage these lipids have higher melting temperatures, due to longer and more saturated acyl chains. Another surprise is that no significant change in sterol content is observed. These results fit within the predominant view that phase separation in membranes requires at least three types of molecules to be present: lipids with high melting temperatures, lipids with low melting temperatures, and sterols.

**SIGNIFICANCE STATEMENT:** When budding yeast shift from growth to quiescence, the membrane of one of their organelles (the vacuole) undergoes liquid-liquid phase separation. What changes in the membrane’s lipids cause this phase transition? Here, we conduct lipidomics of immuno-isolated vacuole membranes. We analyze our data in the context of lipid melting temperatures, inspired by observations that liquid-liquid phase separation in model membranes requires a mixture of lipids with high melting temperatures, lipids with low melting temperatures, and sterols. We find that phase-separated vacuole membranes have higher concentrations of PC lipids, and that those lipids have higher melting temperatures. To conduct our experiments, we developed a tagged version of a protein (Mam3) for immuno-isolation of vacuole membranes.

## INTRODUCTION

During normal growth, cells undergo enormous changes as they adapt to their environment and pass through a sequence of distinct metabolic states (1). For example, when nutrients other than the primary carbon source becomes limiting, *Saccharomyces cerevisiae* (henceforth “yeast”) transition away from fermentative, exponential growth (log stage), through respiratory growth (diauxic shift), to reach a quiescent state (stationary stage) (2). The shift from the logarithmic to the stationary stage is accompanied by striking changes in the yeast vacuole, the functional equivalent of the lysosome in higher eukaryotes. Multiple, small vacuoles fuse so that most cells contain only one large vacuole ((3), and reviewed in (4)), and the vacuole membrane undergoes liquid-liquid phase separation (Fig. 1) (5). As a result, in the stationary stage, the vacuole membrane contains micron-scale domains that are enriched in particular lipids and proteins (6–9). This phase transition is reversible, with a transition temperature roughly 15°C above the yeast’s growth temperature (10). Recent studies suggest that phase-separated domains in vacuole membranes regulate the metabolic response of yeast to nutrient limitation through the TORC1 pathway, which regulates protein synthesis, autophagy, lipophagy, and other processes (8, 9, 11–14).

**Figure 1:**
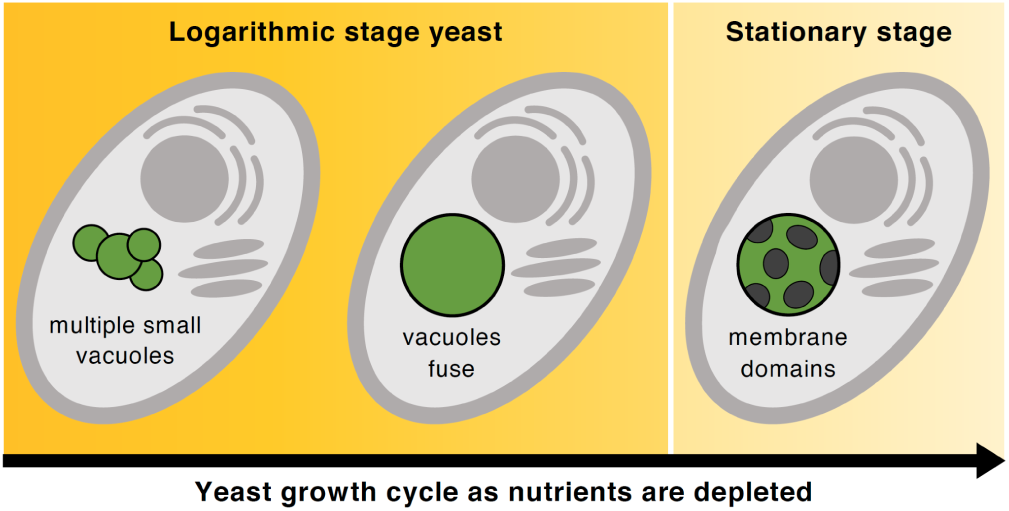
(A) In nutrient-rich media, yeast cultures grow exponentially. In this “logarithmic stage”, each cell contains multiple small vacuoles, the lysosomal organelle of yeast. (B-E) As nutrients become limited, the vacuoles fuse, and yeast enter the “stationary stage”. In this stage, the vacuole membrane phase separates into micron-scale, coexisting liquid phases. Panels C and E are fluorescence micrographs collected as in (10).

An outstanding question in the field has been what the molecular basis is for phase separation in the vacuole membrane. Here, we investigate changes in the lipidome of the vacuole as yeast enter the stationary stage. We expect the lipidome to be important because the few mutations that are known to perturb phase separation of the vacuole involve lipid trafficking and metabolism (8, 9, 15). We expect observable changes in the lipidome because log-stage vacuole membranes do not separate over large shifts in temperature from 30°C to 5°C, whereas stationary stage vacuole membranes do phase separate, even when they are grown over a range of growth temperatures (10). However, we do not necessarily expect the changes driving phase separation to be large. Phase transitions are an effective means of amplifying small signals.

What changes in the lipidome do we expect? In the past, researchers have focused on ergosterol, the predominant sterol in fungi and many protozoans. Multicomponent model membranes containing ergosterol can separate into coexisting liquid phases (16). Moreover, phase separation in isolated vacuoles is reversed through changes in ergosterol levels (8, 10, 17). Klose *et al*. measured whole cell lipidomes through the growth cycle and found that ergosterol decreased from ∼14% of total lipids in the log stage, to ∼10% in the stationary stage (18). However, most ergosterol resides in the plasma membrane and it remained unclear how much of this change could be ascribed to the vacuole membrane alone.

Previous attempts to measure changes in vacuole lipidomes have been limited by the technical challenge of separating vacuole membranes from the membranes of other organelles. This challenge is formidable because vacuole membranes are in contact with other membranes, such as the nuclear envelope, which can co-purify with vacuoles (19, 20). Here, we employ an immuno-isolation technique called MemPrep to efficiently separate vacuole membranes from those of other organelles (21–25). We first identify a bait protein that resides only in the membrane of interest and then genomically fuse a cleavable epitope tag to that bait protein. To achieve roughly equal immuno-isolation efficiencies in the log and stationary stages, we use the membrane protein Mam3 as our bait, which isolates highly enriched membranes from both growth stages at sufficient yields for quantitative lipidomic analyses. This allows us to map the differences in the lipid profiles of the vacuole membrane in the logarithmic and the stationary stages. We then put our lipidomic data into context of existing data of the physical properties of lipids.

## MATERIALS AND METHODS

### Yeast cell culture and microscopy

For lipidomics experiments, we used a *Saccharomyces cerevisiae* strain from the MemPrep library (25) with a bait tag targeted to the C-terminus of Mam3 (a vacuolar membrane protein involved in Mg^2+^ sequestration) (26). The bait tag for immuno-isolation contains a linker region followed by a Myc epitope tag for detection in immunoblotting analysis, a specific cleavage site for the human rhinovirus (HRV) 3C protease for selective elution from the affinity matrix, and three repeats of a FLAG epitope that ensures binding to the affinity matrix. The complete amino acid sequence of the bait tag is: GGGSGGGGSEQKLISEEDLGSGLEVLFQGPGSGDYKDHDGDYKDHDIDYKDDDDK.

From a single colony on a YPD agar plate, 3 ml synthetic complete media was inoculated. Cells were cultivated for 20 hours at 30°C, producing a starter culture. From this starter culture, we inoculated 4 L of synthetic complete media to an optical density at 600 nm (OD_600_) = 0.1 and cultivated the cells at 30°C and 220 rpm constant agitation. For yeast in the log stage, cells were cultivated for approximately 8 hours, until they reached OD_600_ = 1. For yeast in the stationary stage, cells were cultivated for 48 hours. In the stationary stage, roughly 80% of vacuole membranes undergo phase separation into micron-scale domains (8–10, 15).

Yeast were imaged by fluorescence microscopy as described in the supplemental methods section of the Supporting Materials.

### Immuno-isolation of microsomes

Microsomes of vacuole membranes were produced as briefly described in Fig. 2 and as fully described in the supplemental methods of the Supporting Materials. The supernatant of previously prepared magnetic beads (see supplemental methods) was discarded and replaced with 700 µL of fresh IP buffer. Next, 700 µL of the sonicated, crude microsomal vesicle fraction were added to the magnetic beads, for a total volume of 1.4 mL. Vesicles were allowed to bind to the beads (Fig. 2B) by rotating the tubes for 2 h at 4°C in an overhead rotor at 3 rpm to avoid excessive formation of air bubbles.

**Figure 2:**
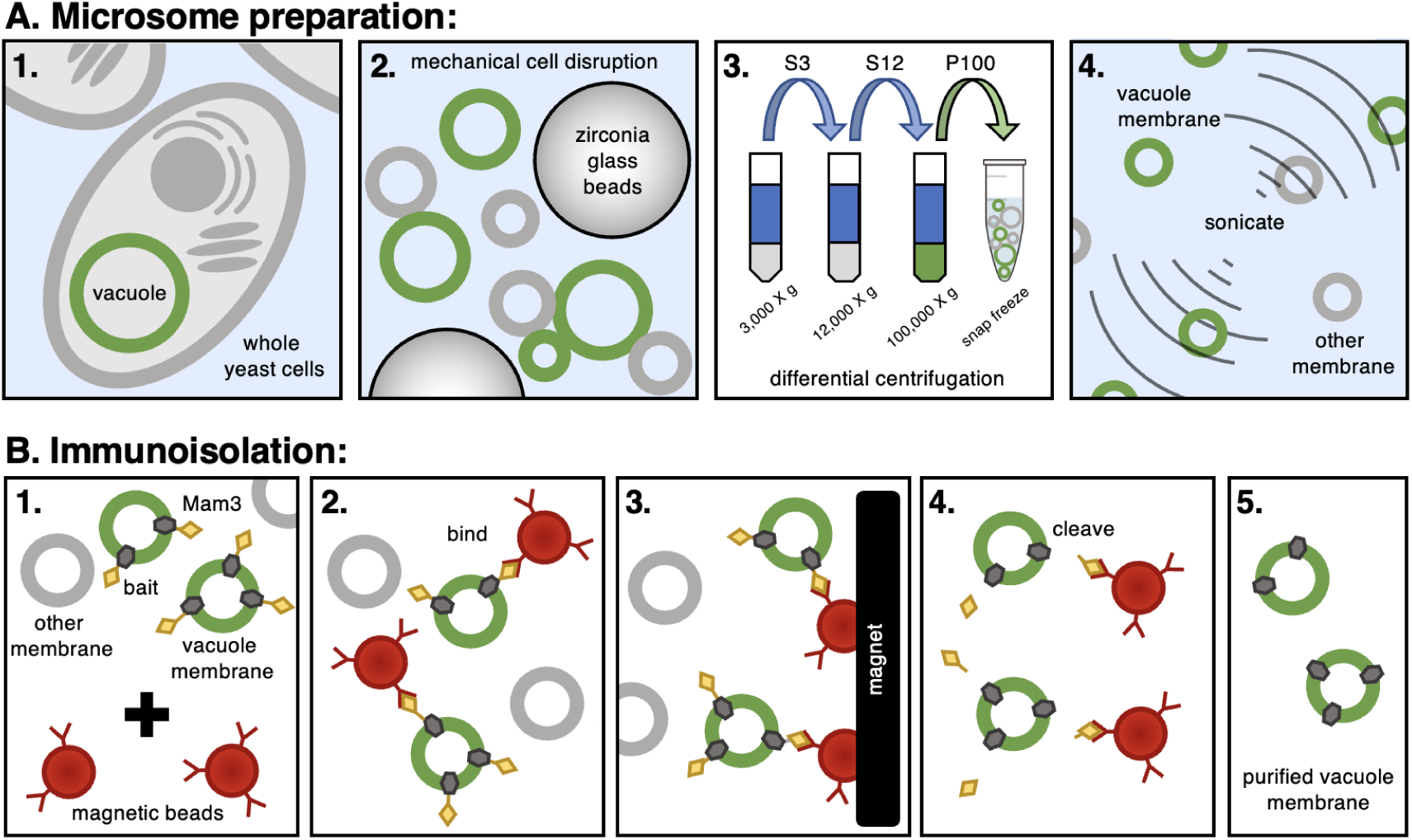
**A.** (**1**) Yeast cells are cultivated to either the log or stationary stage. (**2**) Cells are mechanically fragmented with zirconia glass beads using a FastPrep-24 bead beater. (**3**) A differential centrifugation procedure at 3,234 x g, 12,000 x g, and 100,000 x g is performed to deplete cell debris and membranes from other organelles thereby enriching for vacuole membranes in the microsomal fraction. The fraction containing vacuole membranes (either the supernatant, S, or the pellet, P, is retained. (**4**) Controlled pulses of sonication separates clumps of vesicles and produces smaller microsomes for immuno-isolation (25). Items are not drawn to scale. **B.** (**1**) Top: The microsome solution is enriched in vacuole membranes, which are labeled with a bait tag (myc-3C-3xFLAG) attached to Mam3. Bottom: Microsomes are mixed with magnetic beads coated at sub-saturating densities with a mouse anti-FLAG antibody. (**2**) Antibody-coated, magnetic beads bind to Mam3 in vacuole membranes, but not to other membranes. (**3**) For washing, the affinity matrix (magnetic beads) is immobilized by a magnet, the buffer with all unbound material is removed, and fresh, urea-containing buffer is added. The affinity matrix is serially washed and agitated in the absence of a magnetic field to ensure proper mixing and removal of unbound membrane vesicles. (**4**) Vacuole membranes are cleaved from the beads with affinity purified HRV-3C protease. (**5**) Removal of the magnetic beads leaves purified vacuole membranes for lipidomics.

Magnetic beads, loaded with sample, were collected after a series of washes (Fig. 2B). To this end, the tubes were placed in a magnetic rack and the supernatants were removed. The magnetic beads were washed twice with 1.4 mL of wash buffer (25 mM HEPES, pH 7.0, 1 mM EDTA, 75 mM NaCl, 0.6 M urea), which destabilizes many unspecific protein-protein interactions, and then twice with 1.4 mL IP buffer. After these washes, the magnetic beads were transferred to a fresh 1.5 mL tube and resuspended with 700 µL elution buffer (PBS pH 7.4, 0.5 mM EDTA, 1 mM DTT, and 0.04 mg/mL GST-HRV-3C protease).

To elute vesicles from the affinity matrix (Fig. 2B), the samples were incubated rotating overhead at 3 rpm for 2 h at 4°C with the HRV-3C protease. The tubes were then placed in a magnetic rack to precipitate the magnetic beads, and the supernatant (eluate) containing the purified vacuole membranes was transferred to fresh tubes.

To concentrate the membrane vesicles and for a buffer exchange, the sample was diluted in PBS and transferred to ultracentrifuge tubes. After centrifugation at 264,360 x g for 2 hours at 4°C using a Beckman TLA 100.3 rotor, the supernatant was discarded and the pellet containing purified vacuole membranes was resuspended in 200 µl PBS. Resuspended pellets were transferred to fresh microcentrifuge tubes, snap frozen in liquid nitrogen, and stored at −80°C until lipid extraction and mass spectrometry analysis. Procedures for lipid extraction, acquisition of lipidomics data, and processing of lipidomics data are described in the supplemental methods section of the Supporting Materials.

## RESULTS

### Mam3 is a robust bait protein for immuno-isolation of vacuole membranes

We identified several membrane proteins as candidate “bait proteins” for immuno-isolation of vacuoles in *both* the log stage and the stationary stage of growth. Immuno-isolation with bait proteins provides a high level of organelle selectivity that is not available by standard flotation methods (27, 28). One end of the bait protein (the C-terminus) is equipped with a molecular “bait” (the construct myc-3c-3xFLAG). The bait protein and the membrane in which it resides binds to magnetic beads coated with anti-FLAG antibodies (Fig. 2B). The beads are magnetically immobilized and washed extensively with urea-containing buffers. The isolated membranes are then cleaved from the beads (using the GST-HRV-3C protease) and analyzed by shotgun lipidomics. For our application, a robust bait protein (1) must have its C-terminus available in the cytosol, (2) contain an intramembrane domain anchoring it to the membrane, (3) partition to only the vacuole, and (4) be expressed at high levels in both the log and stationary stage. Initial guesses can be made about whether given proteins are good candidates for bait proteins by consulting the YeastGFP fusion localization database (https://yeastgfp.yeastgenome.org/), and then fusions must be tested in the lab (29, 30).

An obvious candidate for a bait protein was Vph1, which has high expression levels in the log stage (8, 31). Vph1 has been extensively used for visualizing vacuole domains (5, 7–10, 15), and was previously used by us to demonstrate the utility of MemPrep for isolating vacuole membranes in the log stage (25). However, in the stationary stage, expression levels of Vph1 are lower (Fig. S1), resulting in an insufficient yield of membranes by immuno-isolation of the Vph1-bait construct.

In contrast, Mam3 (∼5890 molecules per cell) proved to be an excellent bait protein. We confirmed previous reports that Mam3 localizes to the vacuole membrane (32) by showing that Mam3 colocalizes with FM4-64, a styryl dye that selectively stains vacuole membranes (Fig. S2). In the log stage, when vacuoles do not exhibit domains, Mam3 distributes uniformly on the vacuole membrane. In stationary stage yeast, Mam3 partitions to only one of the two phases of vacuole membranes (Fig. 3 and Fig. S3). Specifically, Mam3 partitions to the same phase as Vph1, which Toulmay & Prinz identified as a liquid disordered (Ld) phase (Fig. S4) (8). Expression levels of Mam3 are equally high in both the log and stationary stages (Fig. 3C). Mam3 outperformed four other candidates for bait proteins (Ypq2, Sna4, Ybt1, and Ybr247C) with the myc-3c-3xFLAG bait tag. We would expect all of these proteins to preferentially partition to the Ld phase.

**Figure 3:**
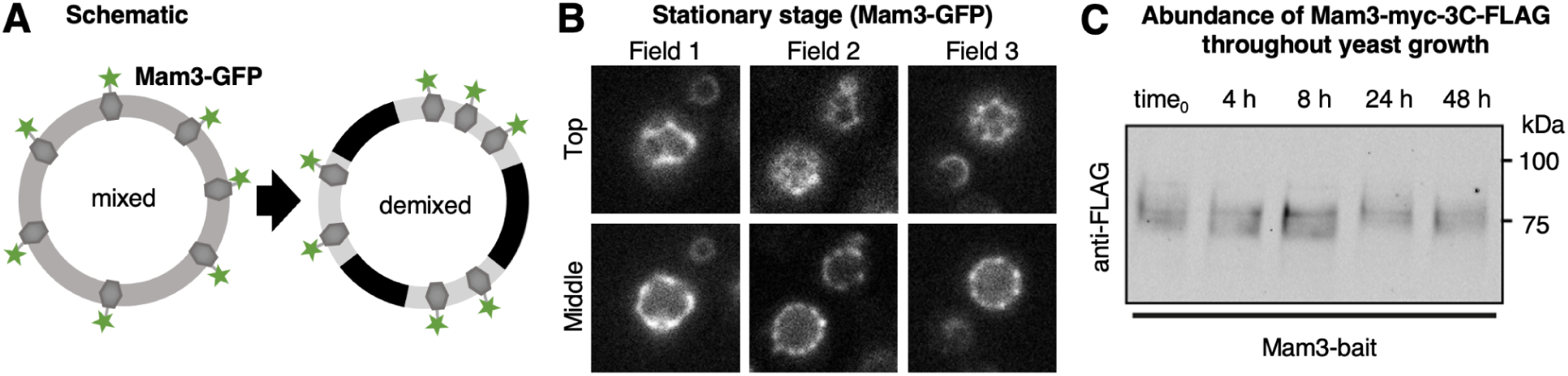
**(A)** In the log stage of growth, lipids and proteins appear uniformly distributed in vacuole membranes of yeast. Mam3, a transmembrane protein in the vacuole membrane, was produced from its endogenous promoter and equipped at its C-terminus either with a bait tag for immuno-isolation or with GFP for fluorescence microscopy. **(B)** *In vivo* fluorescence micrographs of yeast showing that in the stationary stage (after 48 hours of growth), the vacuole membrane phase separates into two liquid phases. Mam3-GFP partitions into only one of these phases (identified as the Ld) phase. Micrographs were taken at room temperature at both the top (“Top”) and the midplane (“Middle”) of vacuoles for each field of view. Wider, representative fields of view are shown in Fig. S3, and corresponding images for Vph1 are in Fig. S4. **(C)** Synthetic complete media was inoculated to an OD600 of 0.1 and cells were harvested by centrifugation after cultivation for 0, 4, 8, 24, and 48 hours. Immunoblot showing robust and stable expression levels of Mam3, visualized using anti-FLAG and fluorescently labeled secondary antibodies, which bind the myc-3c-3xFLAG bait tag on Mam3. Bands for Mam3 are shown in the log (early times) and stationary stage (late times), and positions of molecular weight markers are indicated for reference.

For completeness, we attempted to find complementary bait proteins that preferentially partition to the opposite phase (the liquid ordered, or “Lo” phase) of vacuole membranes in the stationary stage. None were found to be good candidates. For example, the protein Gtr2 partitions to the liquid ordered (“Lo” phase) in yeast vacuole membranes (8). We find that membrane fragments isolated with a bait attached to Gtr2 proteins contain low amounts of proteins known to reside in vacuole membranes (Vph1 and Vac8); the Gtr2 bait protein does not isolate enough vacuole membrane to analyze by lipidomics (Fig. S5). To date, the only other protein known to preferentially partition to the Lo phase of vacuole membranes is Ivy1 (8). Because Ivy1 is an inverted BAR protein rather than a transmembrane protein, it is an unsuitable bait protein.

### Low contamination by non-vacuolar proteins and lipids

A key challenge in isolating pure organellar membranes is that most organelles are in physical contact with other organelles, which can contaminate membrane samples. Most previous isolation attempts have been based on the protocol pioneered by Uchida *et al*. in which yeast are converted into spheroplasts and mildly lysed, and then vacuole membranes are enriched by differential centrifugation and density centrifugation (27, 33, 34) [see Table S1 for comparisons]. Zinser *et al*. noted that lipid droplets “seemed to adhere” to vacuole membranes isolated by this method (7, 28). Contamination of vacuoles by other types of organelles is smaller. When Tuller *et al*. isolated vacuole membranes, they found 0.5% contamination by plasma membranes and 5.5% contamination by cardiolipins, a lipid specific to mitochondria (34). Likewise, Schneiter *et al*. reported that a vacuolar marker protein was enriched ∼15-fold in their vacuolar membrane preparations, with some contamination from the ER and the outer mitochondrial membrane (22).

Here, we couple an immuno-isolation approach with a sonication step that segregates vesicle aggregates and produces smaller microsomes (Fig. 2). We benchmark the purity of isolated vacuole membranes by verifying that protein markers for vacuoles are enriched in post-immuno-isolation fractions and that markers for other organelles are depleted. For example, Dpm1, which localizes to the endoplasmic reticulum, and Por1, which localizes to the mitochondrial outer membrane, are initially present after sonication of microsomes (“load” column in Fig. 4) and are removed following immuno-isolation (“eluate” column). In contrast, Vac8 and Vph1, which localize to the vacuole, are enriched in the immunoisolation eluate. The Mam3 bait protein cannot be detected with anti-FLAG antibodies after elution, because the FLAG epitope is cleaved off to release vacuole membranes from the magnetic beads.

**Figure 4:**
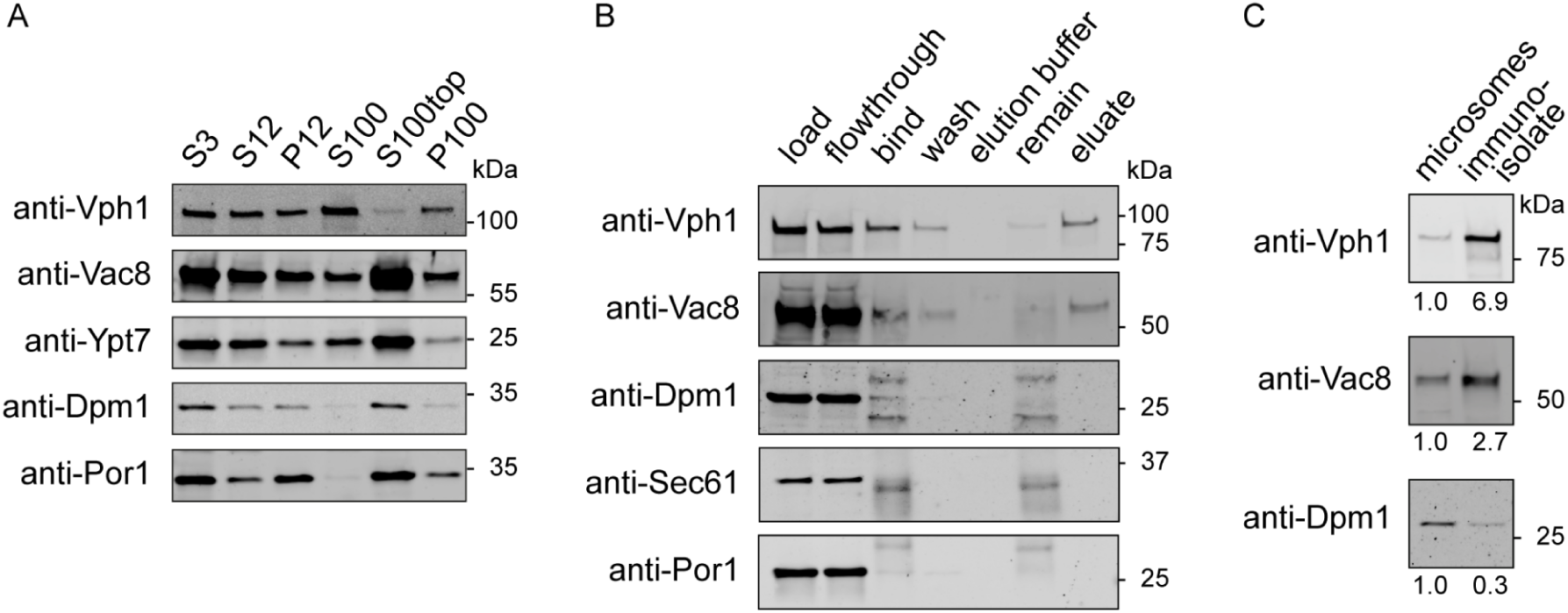
Immuno-isolation from cells expressing the Mam3-bait construct. **(A)** After differential centrifugation, proteins that reside in membranes of the endoplasmic reticulum (Dpm1) and mitochondria (Por1) are present in microsome preparations of yeast (“P100” column), as reported by antibodies to those proteins (anti-Dpm1 and anti-Por1). **(B)** P100 microsomes are sonicated and used as input for immuno-isolation (load). Bands for vacuole markers (Vph1 and Vac8) show highly specific binding to antibody-coated magnetic beads (“bind” column), mitigating substantial loss in the flow through step (“flowthrough” column). The anti-Dpm1 immunoblot shows two additional bands in the fractions containing magnetic beads (in the “bind” and “remain” columns) originating from the FLAG antibody light chain (∼25 kDa) and the coating of protein G (∼37 kDa). Protein G also leads to a band in the anti-Sec61 and anti-Por1 immunoblots. Immuno-isolation removes membranes of the endoplasmic reticulum and mitochondria (seen by an absence of Dpm1, Sec61, and Por1 in the “eluate” column) and retains only vacuole membranes for lipidomics (seen by bands for Vac8 and Vph1 in the “eluate” column). **(C)** Equal amounts of total protein (0.45 µg) were loaded to quantify enrichment over crude microsomes of organelle markers from immunoblot signals.

We also evaluated the purity of isolated vacuole membranes by assessing contamination by lipids known to be in other organelles. We find that only ∼1% of lipids in isolated membranes (0.8 ± 0.4%) contain cardiolipins from mitochondria. Similarly, triacylglycerols (TAG) and ergosterol esters are “storage lipids” found in the hydrophobic core of lipid droplets (9). For yeast in the log stage, we find that only ∼2.5% of all lipids of immuno-isolated vacuoles are TAG, and < 1% are ergosterol esters (Fig. S6). In comparison, in whole cell extracts of the same cells, we find 9% TAG and 5% ergosterol esters (Fig. S7), in agreement with literature values of 10 ± 1% TAG for equivalent yeast and conditions (35). A characteristic feature of vacuole membranes in the log stage of growth is the almost complete absence of phosphatidic acid (PA) (25), which we confirm for vacuole membranes isolated using the Mam3 bait (Fig. 5A).

**Figure 5:**
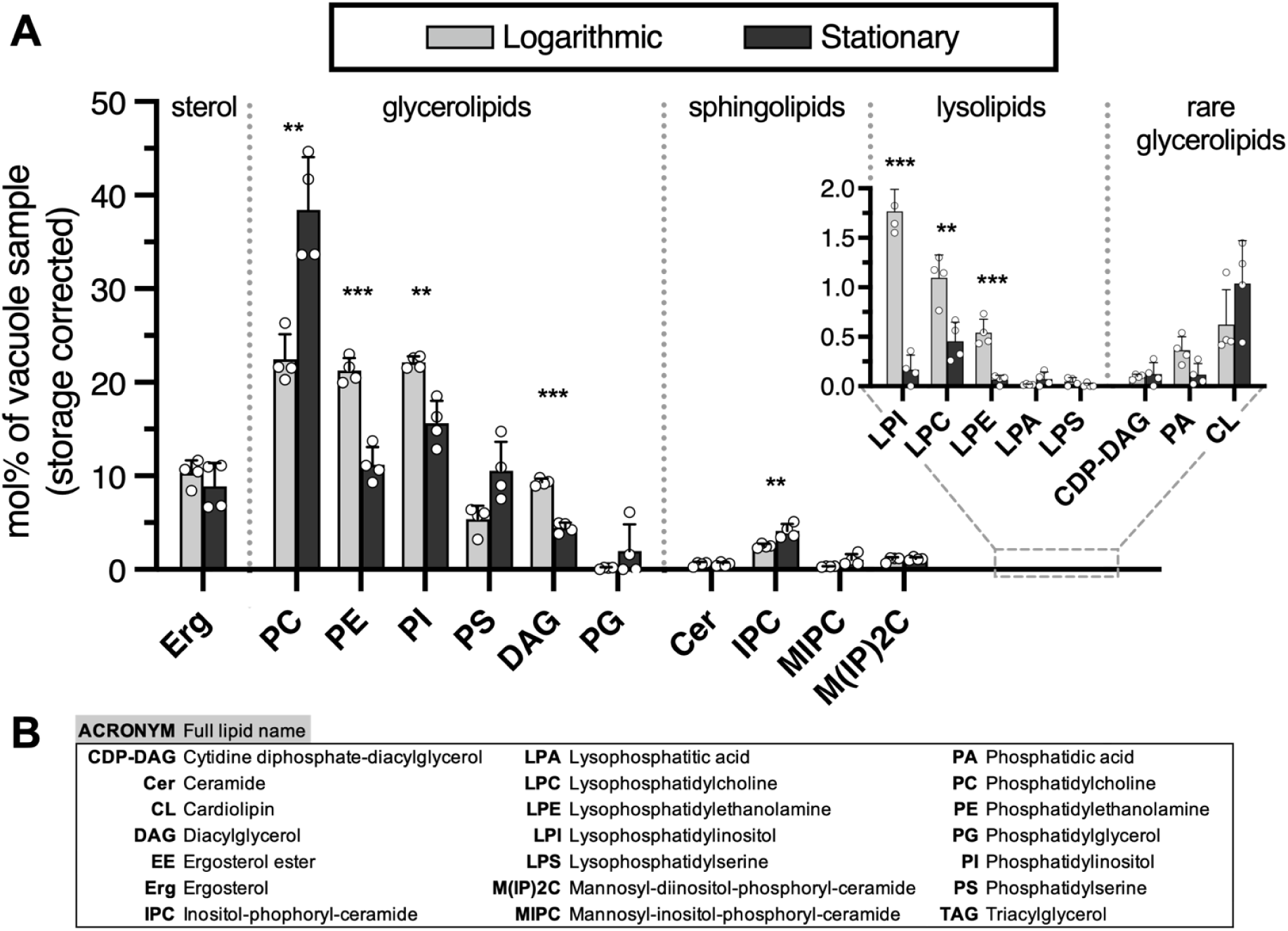
**(A)** Abundances of lipids in yeast vacuole membranes in the logarithmic and stationary stages of growth. A large increase is observed in PC lipids. Concomitant decreases are observed in PE-lipids, PI-lipids, and DAG. Error bars are standard deviations of four vacuole samples immuno-isolated on different days; individual data points are shown by white circles. Data exclude storage lipids (ergosterol ester and triacylglycerol), which are attributed to lipid droplets. Corresponding graphs that include storage lipids are in Fig. S6 and full data sets are in the supporting information. Statistical significance was tested by multiple t-tests correcting for multiple comparisons (method of Benjamini et al. (42)), with a false discovery rate Q = 1%, without assuming consistent standard deviations. *p < 0.05, **p < 0.01, ***p < 0.001. **(B)** Acronyms of lipid types.

Isolation of pure vacuole membranes is particularly challenging in the stationary stage, when lipid droplets are produced in high numbers and are in intimate contact with vacuole membranes (7, 9, 15, 36, 37). Nevertheless, we still find low contamination. For immuno-isolated vacuoles from yeast in the stationary stage, we find only ∼13% TAG and ∼0.5% ergosterol esters (Fig. S6). In comparison, in whole cell extracts from the same cells, we find three times more TAG (>35%) and an order of magnitude more ergosterol esters (>6%) (Fig. S7). High levels of ergosterol esters persist in vacuoles separated by density gradient methods from log-stage yeast grown in YPD media (28), and may be even higher for equivalent yeast grown in synthetic complete media (35). These data highlight the value of isolating vacuole membranes by immuno-isolation. In the sections below, we exclude storage lipids because they likely originate from lipid droplets and because their solubility is low (< 3% for TAG) in glycerolipid membranes and likely even lower in membranes containing sterols (38, 39).

Breaking contact sites between organelles results in highly purity immuno-isolated membranes. However, like every technique, it has tradeoffs. As noted by Zinser and Daum, “rupture of intact vacuoles may liberate proteases” (40). Therefore, we perform all steps of MemPrep at 4°C; every increase in temperature of 10°C doubles the rate at which proteases degrade proteins. This constraint prevents us from isolating membranes from the same population of vacuoles at two temperatures, a low temperature at which the membrane phase separates, and a high temperature at which the membrane is uniform.

### Logarithmic stage lipidomes have high levels of PC, PE, and PI lipids

We quantified the lipidome for ∼520 individual lipid species, in ∼20 lipid classes. The list of relevant lipids is shorter in yeast than in many other cell types because yeast glycerolipids typically contain at most two unsaturated bonds, one in each chain (41). In general, the PC, PE, and PI lipid classes each constitute ∼20% of the log-stage vacuole lipidome (Fig. 5). Lipidomes of log-stage vacuole membranes immuno-isolated via MemPrep using Mam3 as a bait are in excellent agreement with those we previously isolated with a Vph1 bait protein (Fig. S8) (25). It is difficult to assess whether these lipidomes agree with data determined by more traditional methods. Although Zinser *et al*. (28) reported that levels of PC lipids were 2x the level of PE or PI lipids in log-stage vacuoles, it is unclear if the yeast they used were grown to early log stage or a later stage. Whole-cell lipidomes by Reinhard et al. (35) imply that the difference cannot be simply ascribed to the use of YPD media by Zinser and coworkers versus synthetic complete media.

The importance of isolating vacuoles from whole-cell mixtures is reflected in differences between their lipidomes. For log-stage cells grown in synthetic complete media, the fraction of IPC and PA lipids are roughly 2 and 10 times greater in the whole cell, respectively, whereas the fraction of DAG lipids is roughly 2 times higher in the vacuole than the whole cell (18, 35).

### Phase separation in vacuole membranes is not likely due to an increase in ergosterol

A central question is why log-stage vacuole membranes do not phase separate, whereas stationary stage membranes do. Ergosterol is the major sterol in yeast, and it has been suggested that phase separation of vacuole membranes could be due to an increase in ergosterol (8). An increase in ergosterol would be consistent with a report that the ratio of filipin staining of sterols in vacuole versus plasma membranes is higher in the stationary stage than the log stage, although vacuole and plasma membrane levels were not measured independently (10, 15). It would also be consistent with the expectation that esterified sterols in lipid droplets are mobilized in the stationary stage by lipophagy, which is required for the maintenance of phase separation in the vacuole membrane (17, 21, 33, 37). Ergosterol levels in the vacuole membrane do not necessarily reflect levels in the whole cell, which includes lipid droplets. We find that the mole percent of ergosterol in whole cell extracts remains constant at ∼10% from the log stage to the stationary stage (Fig. S7); Klose et al. previously reported a decrease from ∼14% to ∼10% in the early stationary phase (18).

We find statistically equal fractions of ergosterol in vacuole membranes in the log and stationary stages: 10 ± 1% in the log stage and 9 ± 2% in the stationary stage (Fig. 5). Uncertainties represent standard deviations for four independent experiments of each type, and the data exclude storage lipids of ergosterol esters and TAGs. These levels of sterol are often sufficient for separation of model membranes into two liquid phases (43–49). Higher levels are not necessarily better at promoting coexisting liquid phases; the solubility limit of ergosterol in membranes of lipids with an average of one unsaturation is 25-35% (50–53) and solubility may be lower when lipid unsaturation is higher, as in biological membranes.

We cannot rule out that immunoisolation with Mam3 under-samples the liquid ordered phase (which contains higher concentrations of ergosterol per unit area than the liquid disordered phase, based on brighter staining by filipin (8)). Nevertheless, our conclusion that vacuole phase separation is not likely driven by an increase in ergosterol is consistent with other results in yeast; we previously found that log-stage vacuole membranes phase separate upon depletion (rather than addition) of ergosterol (10). Other results in the literature are more difficult to interpret. It is not known whether ergosterol and other lipids move into yeast vacuoles or out of them when sterol synthesis and transport are impaired (9, 11, 12, 15), when lipid droplets are perturbed (9, 13), or when drugs are applied to manipulate sterols (8, 17) (all of which can disrupt the formation or maintenance of vacuole membrane domains).

Results in other cell types are not necessarily applicable to yeast. As in yeast vacuoles, depletion of sterol from modified Chinese hamster ovary cells results in plasma membrane domains, and recovery of photobleached lipids is consistent with the domains and the surrounding membrane both being fluid (54). Similarly, depletion of sterol from giant plasma membrane vesicles (GPMVs) of NIH 3T3 fibroblasts causes phase separation to persist to higher temperatures (55). Membrane domains in other types of sterol-depleted cells (e.g. (56)) may also prove to be due to phase separation, although it is always important to verify that cells are still living (as in (56)) and to identify when membrane properties such as lipid diffusion, dye partitioning, domain shape, and/or domain coalescence reflect liquid rather than solid phases (as in (57–59)). Different results are observed in vesicles derived from other cell types. Depletion of sterol from GPMVs of RBL cells restricts phase separation to lower temperatures (60). Similarly, when phase separation is restricted to lower temperatures in zebrafish GPMVs (because the cells are grown at lower temperature), sterol levels are lower (61). Results from these different cell types are not in conflict; addition and depletion of sterol have been suggested to drive membranes toward opposite ends of tie-lines (away from phase separation and toward a single, uniform phase), based on observations in GPMVs of RBL-2H3 cells (62), CH27 cells (63), and model membranes (16, 43, 55, 64).

### Changes in vacuole lipidome from the log stage to the stationary stage

If phase separation in vacuole membranes is not due to an increase (or even a change) in the fraction of ergosterol, could it be due to changes in other lipids? In the remainder of this paper, we explore how the vacuole lipidome changes from the log stage to the stationary stage, starting with Fig. 5. A first impression from Fig. 5 is that the lipid populations “are more alike than they are different” (to quote Burns et al. (61), who compared lipidomes from zebrafish GPMVs that phase separate to ones that do not). It makes sense that the two populations are similar, because the cells from which they are extracted are separated by only two days of cultivation.

Nevertheless, significant changes in the lipidomes are apparent. The most dramatic change is the increase in the fraction of PC lipids. This jump is notable because PCs are a large fraction of the vacuole’s lipids and because the change does not occur on a whole-cell level (18). In contrast, decreases in PE, PI, and DAG lipids mirror decreases in the whole-cell lipidome.

Evaluating the lipidome for each lipid type is essential because average values can obscure shifts. For example, the average number of double bonds per lipid chain in vacuole membranes for all lipids (excluding storage lipids of TAG and EE) remains constant from the log stage (0.67 ± 0.07) to the stationary stage (0.65 ± 0.06). Even when the data are restricted to glycerolipids with two chains, only a small decrease in lipid unsaturation is apparent (Fig. 6A). In contrast, when phosphatidylcholine lipids are analyzed on their own (because there is a large increase in PC lipids from the log to the stationary stage), a large change in unsaturation appears (Fig. 6C). The fraction of PC lipids that contain two unsaturated acyl chains significantly decreases from 83 ± 3% in the log stage to 54 ± 2% in the stationary stage. A concomitant increase in lipids with only one unsaturated chain occurs, from 16 ± 3% in the log-stage vacuoles to 44 ± 2% in the stationary stage vacuoles (Fig. 6C). Similarly, the average length per lipid chain for all lipids in the vacuole (excluding TAG and EE) remains constant from the log stage (16.6 ± 0.8 carbons) to the stationary stage (16.9 ± 1.8 carbons), whereas by restricting the data to glycerolipids with two chains (Fig. 6B) or PC lipids (Fig. 6D), it is clear that chain lengths slightly increase for these lipids, largely due to a decrease in lipids with a total acyl chain length of 32 carbons, in favor of longer lipids.

**Figure 6:**
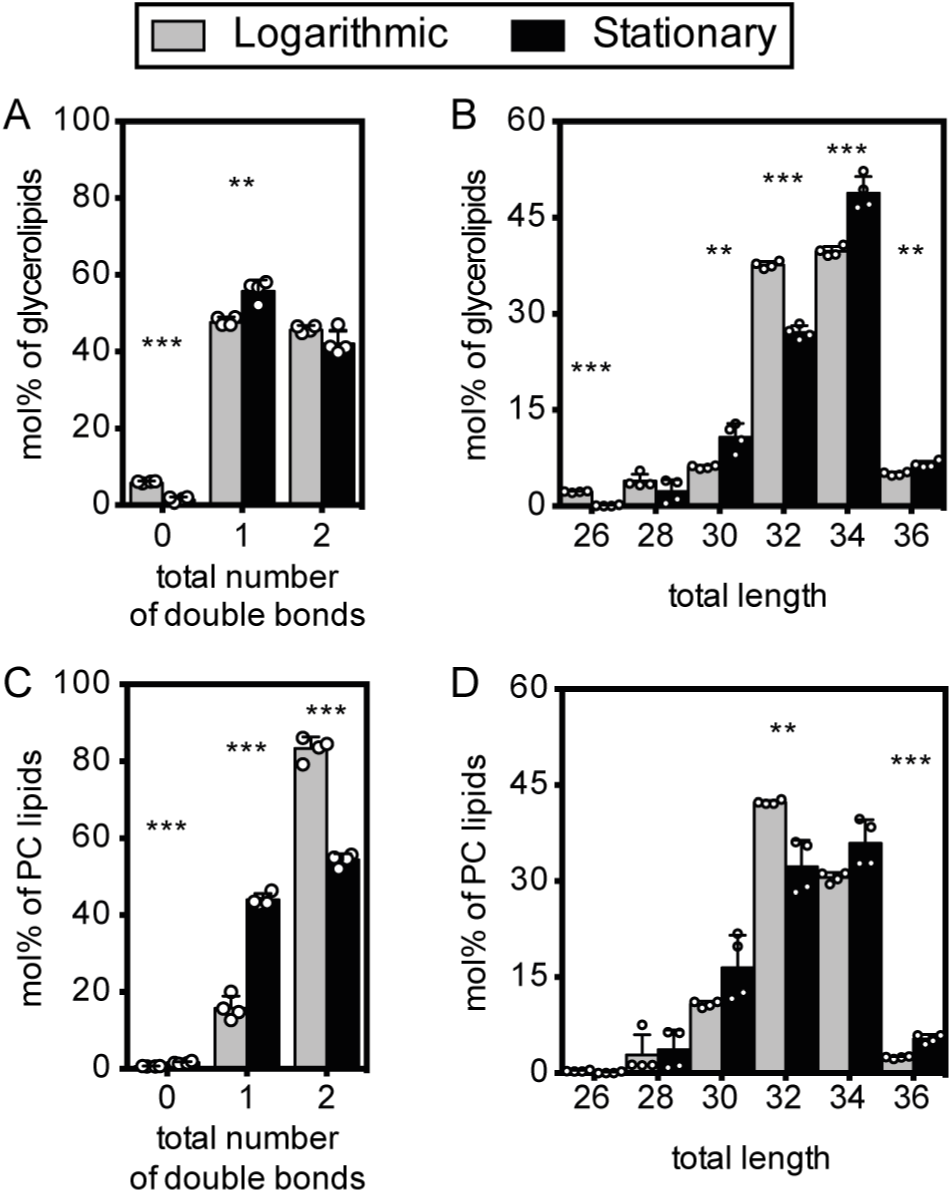
Molecular features of glycerolipids and phosphatidylcholine (PC) in vacuole membranes. (A) Total number of double bonds in glycerolipids that have two acyl chains (CDP-DAG, DAG, PA, PC, PE, PG, PI, PS) in the log (gray) and stationary (black) stages. These lipids constitute 81 ± 2% and 82 ± 4% of all membrane lipids in the log and stationary stage vacuole membrane, respectively. (B) Total length of acyl chains in glycerolipids with two chains. A shift from the log stage to the stationary stage is accompanied by a small increase in lipid lengths. (C) Total number of double bonds in phosphatidylcholine lipids (PC). Number of double bonds in PC lipids. A shift from the log to the stationary stage is accompanied by a large decrease in the proportion of PC lipids with two unsaturated acyl chains. (D) Total length of acyl chains in PC lipids. Statistical significance was tested by multiple t-tests correcting for multiple comparisons (method of Benjamini et al. (42)), with a false discovery rate Q = 1%, without assuming consistent standard deviations. *p < 0.05, **p < 0.01, ***p < 0.001.

### PC lipids shift to higher melting temperatures in the stationary stage

PC lipids are abundant. Like PE and PI, this lipid class constitutes a large fraction (20-25%) of all lipids in log stage vacuole membranes. When yeast transition to the stationary stage, the proportion of PC lipids shoots up to ∼40% of vacuolar lipids, the largest increase of any lipid type. Therefore, it seems likely that the onset of phase-separation of vacuole membranes is linked to changes in the fraction of PC lipids and their acyl chain compositions. This result is consistent with the observation that yeast knockouts of *OPI3* and *CHO2* (which diminish synthesis of PC lipids (65, 66)) exhibit a smaller proportion of vacuoles with domains (8). The mole fractions of every PC lipid in log and stationary stage vacuoles is provided in Data File S1.

To gain intuition about membrane properties imparted by the diverse set of PC lipids, we mapped the lipidomics data onto a physical parameter, the temperature at which each lipid melts from the gel to the fluid phase (*T*_melt_). Melting temperatures are related to phase separation. Model membranes phase separate into micron-scale domains when at least three types of lipids are present: a sterol, a lipid with a high *T*_melt_, and a lipid with a low *T*_melt_ (43, 44, 46–48, 64, 67). In turn, lipid melting temperatures are affected by the lipid’s headgroup, chain length, and chain unsaturation (68–70). When only one of the lipids in a model membrane is varied, a linear relationship can result between the lipid’s *T*_melt_ and the temperature at which the membrane demixes into liquid phases (43, 71). Of course, yeast vacuole membranes experience changes in more than one lipid type and in the relative fractions of lipid headgroups, which likely breaks simple relationships between *T*_melt_ of lipids and the membrane’s mixing temperature (43, 44, 72). However, we are unaware of any single physical parameter that is more relevant than *T*_melt_ for characterizing lipid mixtures that phase separate.

The formidable task of compiling *T*_melt_ values of all PC lipids is less onerous for yeast vacuoles than for other cell types because each carbon chain of a yeast glycerolipid has a maximum of one double bond (41). This fact provides an easy way to evaluate measurement uncertainties because nonzero entries for polyunsaturated phospholipids in Data File S1 (typically well below 0.2%) must be due to error in the process of assigning lipid identities to mass spectrometry data or are due to fatty acids from the medium. Some *T*_melt_ values are available in the literature (68, 70, 73–75). For many others, we estimated *T*_melt_ values from experimental trends (Table S2-S4 and Fig. S9). This procedure yielded *T*_melt_ values for 96% of PC lipids in log-stage vacuoles, and 94% in stationary stage vacuoles.

By compiling all of our data on PC lipids (Fig. 7), we find that they undergo significant acyl chain remodeling from the log stage to the stationary stage in terms of their melting temperatures. Values of *T*_melt_ are much higher for vacuole PC lipids in the stationary stage compared to the log stage (the weighted average *T*_melt_ is −31°C in the log stage and −21°C in the stationary stage). Graphically, Fig. 7A shows this shift as an increase in the mole percent of lipids with high *T*_melt_ (dark bands).

**Figure 7:**
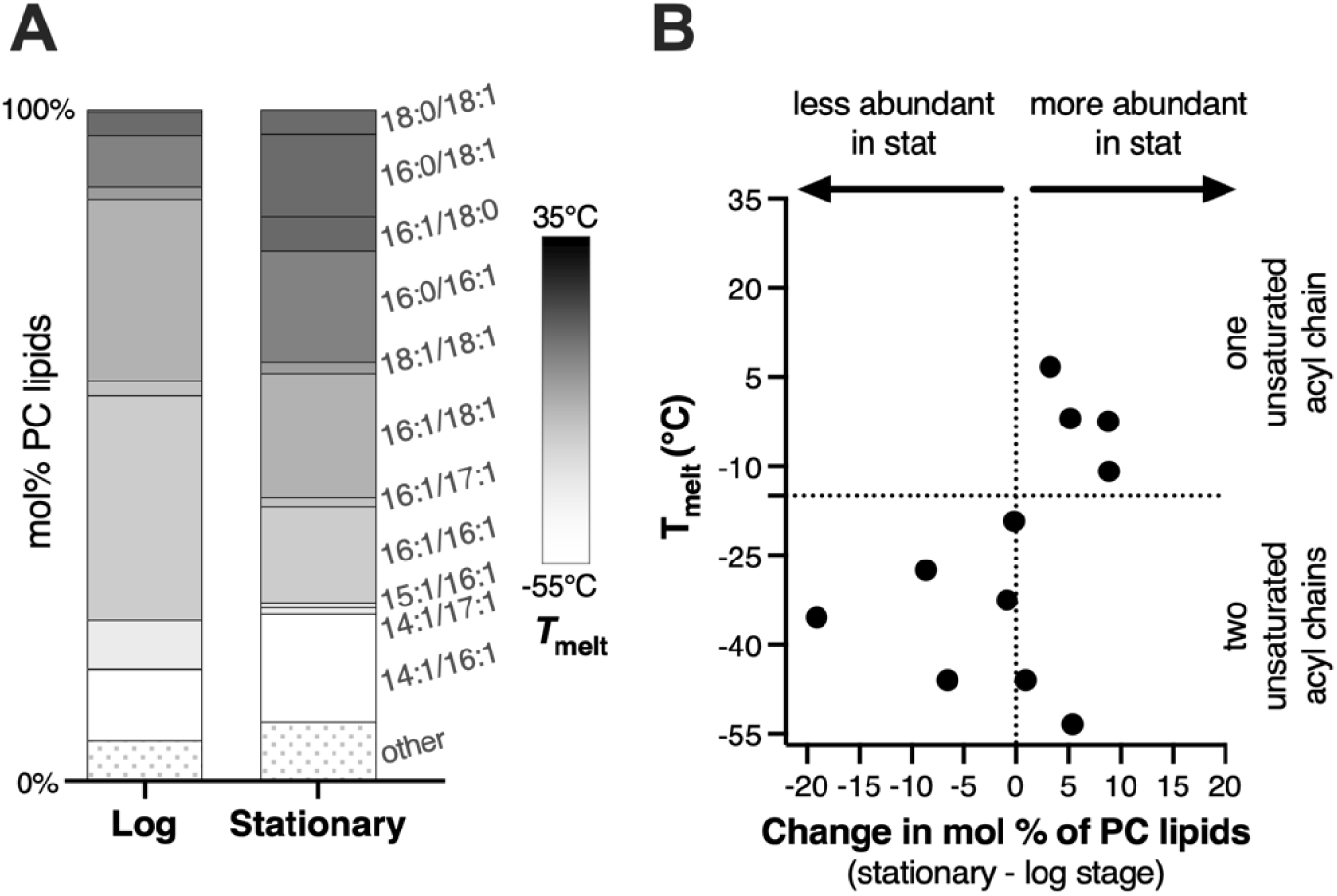
Melting temperatures of PC lipids in vacuole membranes. (A) *T*_melt_ values (represented by a grayscale) are higher for PC lipid in the stationary stage than the log stage. Lipids contributing less than 1 mol% are categorized as “other”. (B) From the log to the stationary stage, there is an overall loss of PC lipids with low values of *T*_melt_ (lower left quadrant) and a gain of PC lipids with high values of *T*_melt_ (upper right quadrant). Fig. S10 presents an alternative way of plotting these data.

The increase in *T*_melt_ correlates with an increase in lipid saturation. In the shift from the log to the stationary stage, PC lipids with two unsaturated chains (lower *T*_melt_) become less abundant, and PC’s with one unsaturated chain (higher *T*_melt_) become more abundant (Fig. 7B and Fig. S10). This large increase in saturation of PC lipids is not reflected in all glycerolipids of the vacuole (Fig. S11).

The increase in average *T*_melt_ of PC lipids that we observe is robust to any possible oversampling of the Ld phase by the Mam3 immunoisolation procedure. Because Lo phases typically contain higher fractions of lipids with higher melting temperatures and orientational order (43, 76) sampling more of the Lo phase would be expected to further increase average *T*_melt_ values.

The spread in the distribution of lipid melting temperatures is likely to be as important as the average. This is because phase separation persists to higher temperatures in model membranes when the highest *T*_melt_ is increased (43, 72) and when the lowest *T*_melt_ is decreased (71), at least when those lipids have PC headgroups and do not have methylated tails. These results can be put into broader context that the addition of any molecule that partitions strongly to only one phase (e.g. a high-*T*_melt_ lipid to the Lo phase and a low-*T*_melt_ lipid to the Ld phase) should lower the free energy required for phase separation (77–79).

In Fig. 7A, PC lipids in log-stage vacuole membranes cluster around intermediate values of *T*_melt_. In the stationary stage, some lipids are replaced by lipids with higher values of *T*_melt_, and some are replaced by lipids with lower values. If the vacuole membrane contained only PC lipids, we would conclude that those membranes would be more likely to phase separate in the stationary stage, because their overall melting temperatures are higher and because the distribution of melting temperatures is broader.

### PE lipids have similar melting temperatures in log and stationary stage

We applied the same procedure to finding melting temperatures of the PE lipids in yeast vacuoles, compiling values in the literature (68, 69, 80, 81) or estimating them (Tables S2-S3 and Fig. S12). PE lipids are particularly relevant because they undergo the largest decrease (from ∼20% to ∼10%) of all lipids in yeast vacuoles. This is interesting because high levels of PE lipids are crucial for maintaining fluid membranes in insect cells, which are devoid of sterol synthesis (82).

In contrast to PC lipids, the acyl chain composition of PE lipids is not heavily remodeled from the log stage to the stationary stage. This result is seen as a clustering of data points along the vertical dashed line in Fig. 8B. Accordingly, the melting temperatures of PE lipids do not change significantly: the weighted average *T*_melt_ is −7°C and −5°C in the log and stationary stages, respectively. Similarly, by eye from Fig. 8A, the distribution of *T*_melt_ values for PE lipids are similar in the log stage and the stationary stage.

**Figure 8:**
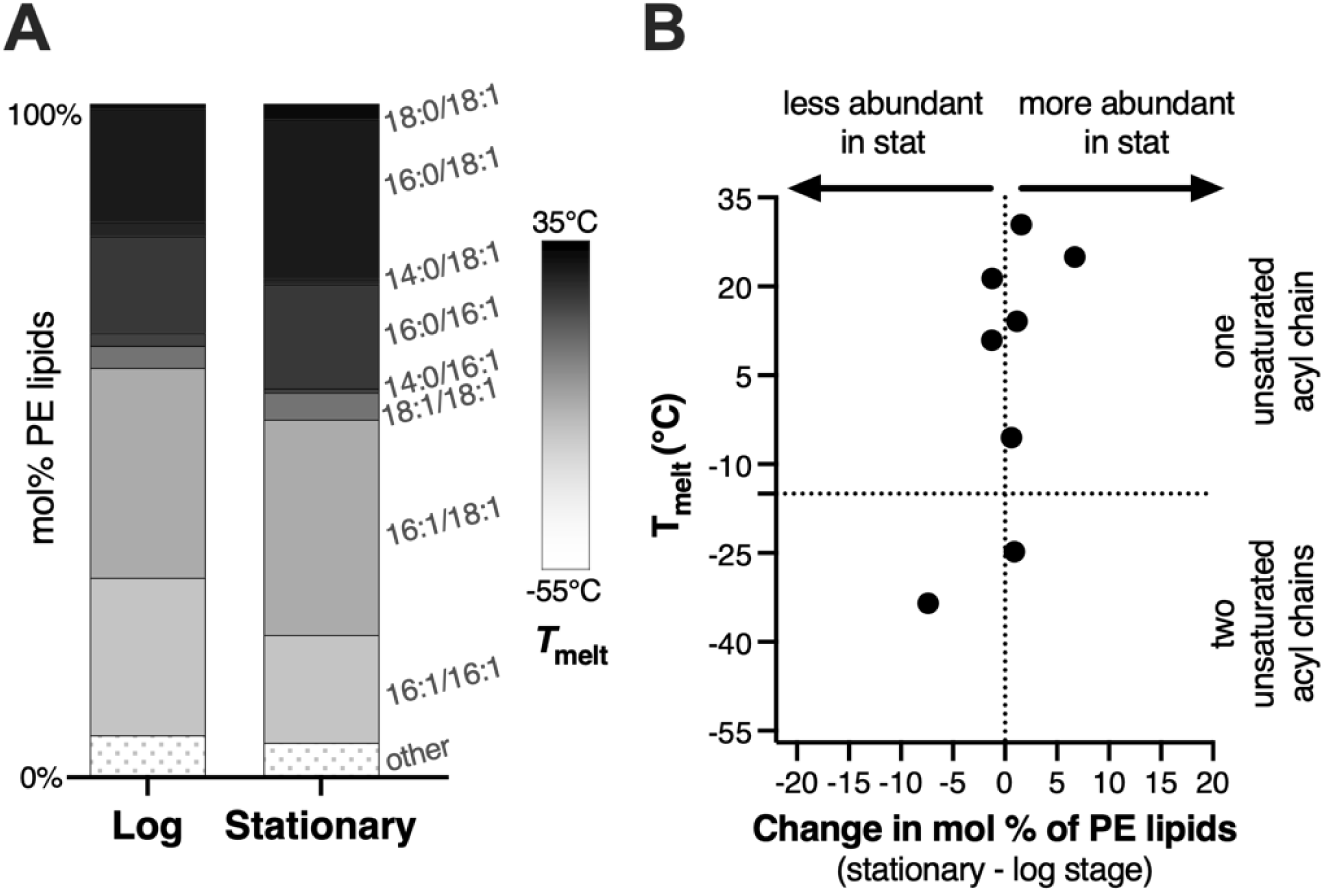
Melting temperatures of PE-lipids in vacuole membranes. (A) *T*_melt_ values (represented by a grayscale) are similar for vacuole PE-lipids in the stationary stage and the log stage. Lipids contributing less than 1 mol% are categorized as “other”. (B) Changes in mol% of PE-lipids (x-axis) with each melting temperature (y-axis) are small; data cluster along the vertical line for zero change in mol%.

### Trends in *T*_melt_ for all glycerolipids

Literature values for lipid melting temperatures are not available for all lipid types, so plots like Fig. 7 and 8 cannot be reproduced for all headgroups. The next best option is to apply known trends of how lipid chain length and saturation affects lipid *T*_melt_ ((68, 70) and Table S3). We estimate changes in *T*_melt_ from the log to the stationary stage for each lipid headgroup separately because it is not clear that combining data sets for all lipids would yield insight about whether a membrane would be more likely to phase separate. For example, two lipids can have the same melting temperature, but membranes containing those lipids can phase separate at different temperatures. For example, pure bilayers of palmitoyl sphingomyelin, 16:0SM, and dipalmitoyl phosphatidylcholine, di(16:0)PC, have the same *T*_melt_ (68, 83), but multi-component model membranes containing 16:0SM phase separate at a higher temperature than membranes containing di(16:0)PC (43, 44), possibly because sphingomyelins have additional opportunities to hydrogen bond with the sterol (84, 85). Similarly, even though PE lipids have higher melting temperatures than equivalent PC lipids (Fig. S13), sterol (cholesterol) partitioning is lower in PE membranes than in PC membranes (86), implying that interactions between sterols and PE lipids are less favorable.

Changes in lipid saturation have much larger effects on melting temperatures than changes in chain length. We find that increasing the average, summed length of both lipid chains by one carbon results in an increase in *T*_melt_ of 3.6°C for PC and PE lipids (because increasing the total length from 32 to 34 carbons results in an increase in *T*_melt_ of 7.2°C, as in Table S5). Increasing the saturation of the lipid by one bond results in an increase in *T*_melt_ of 30.3°C. Therefore, an increase in *T*_melt_ of, say, 5°C can be achieved either by increasing the average total chain length by 5/3.6 = 1.4 carbons or by increasing the average saturation by 5/30.3 = 0.17 bonds (dashed line in Fig. 9).

**Figure 9:**
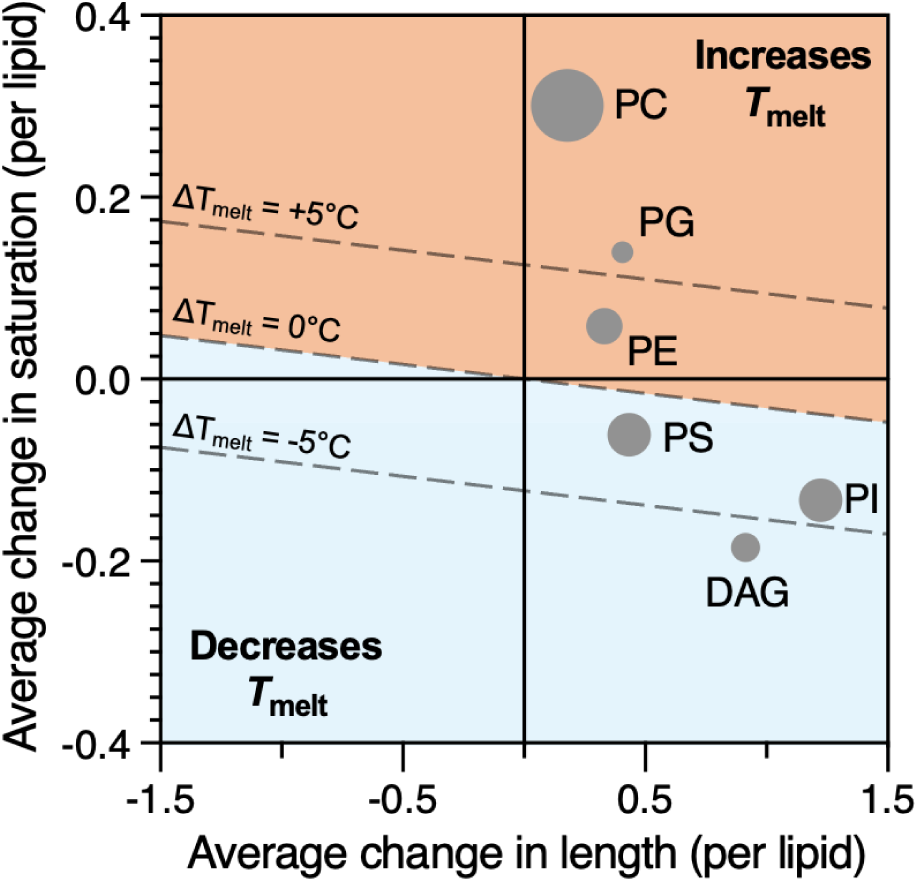
At the origin, there is no change in average lipid length (x-axis) and no change in average lipid saturation (y-axis). Each point represents a different glycerolipid species, and each lipid experiences a change in both length and saturation. The size of each point roughly represents the relative abundance of that lipid class in the stationary stage. Changes in acyl chain saturation have large effects on *T*_melt_ compared to changes in length. Lipids located above the diagonal line of “Δ*T*_melt_ = 0” are expected to experience an increase *T*_melt_. Those below the line are expected to experience a decrease.

We know how chain length and saturation changes from the log stage to the stationary stage for all abundant glycerolipids in yeast vacuoles (PC, PE, PI, PS, PG, and DAG) (Fig. 9 and Fig. S11). By assuming that the melting temperatures of these lipids follow similar trends, we find that *T*_melt_ likely increases for most lipid types during the shift from log to stationary stage. In Fig. 9, changes in chain length or saturation that result in an increase in *T*_melt_ fall within the shaded region toward the top-right of the graph, and the area of each symbol represents the fraction of the lipid type in the stationary stage. The largest increase in *T*_melt_ is for PC lipids, which are the most abundant lipids in the stationary stage. Lipids for which *T*_melt_ likely decreases (PG and DAG) have relatively low abundance. Increases in lipid saturation (and, hence, increases in lipid *T*_melt_ and orientational order) have previously been correlated with phase separation persisting to higher temperatures in GPMVs of zebrafish cells (61).

An alternative mechanism by which eukaryotic cells may influence lipid melting temperatures is through the introduction of highly asymmetric chains. Burns et al. find that zebrafish GPMVs with lower transition temperatures have more highly asymmetric lipids (61). Similarly, when *S. japonicus* fission yeast cannot produce unsaturated lipids (due to anoxic environments), they increase the fraction of lipids with asymmetric acyl tails, which may maintain membrane fluidity (87). However, some yeast (e.g. *S. pombe*) appear to be biochemically unable to use this strategy (87). Although highly asymmetric lipids can appear in *S. cerevisiae* membranes (35), here we find no significant population or change in highly asymmetric lipids vacuole membranes from the log stage to the stationary stage (Table S6).

### Addition of ethanol to isolated, log-stage vacuoles does not cause membrane phase separation

Yeast membranes undergo many changes from the log to the stationary stage that are not captured by lipidomics. For example, as yeast consume glucose, they produce ethanol, some of which partitions into yeast membranes. In model membranes, addition of ethanol results in disordering of lipid acyl chains (88). Lipidomics does not quantify the percent ethanol in membranes. Nevertheless, we can assess whether ethanol on its own is sufficient to cause log-stage vacuole membranes to phase separate. Here, yeast were grown at 30°C in synthetic complete media (with 4% glucose) for < 24 hours until the culture reached an optical density of 0.984. Vacuoles were isolated as in (5), incubated on ice in either 10% v/v ethanol for 1.5 hours or 20% v/v ethanol for 2 hours, and then imaged at room temperature. Neither population of vacuoles showed evidence of phase separation; only one vacuole showed clear evidence of membrane phase separation, which could not be attributed to the addition of ethanol.

The result that ethanol, on its own, is not sufficient to cause log-stage vacuole membranes to phase separate is consistent with literature reports. Although ethanol enhances phase separation in vacuole membranes in model membranes (by increasing the temperature at which coexisting Lo and Ld phases persist) (79), it suppresses phase separation in giant plasma membrane vesicles from rat basal leukemia cells, which have lipid compositions more similar to yeast vacuole membranes (78).

## DISCUSSION

This special issue is dedicated to Klaus Gawrisch, who directed the first experiments to quantify ratios of lipids in Lo and Ld phases of ternary model membranes with high accuracy (76). The resulting NMR data showed that Lo and Ld phases differed primarily in their fractions of phospholipids with high and low melting temperatures (rather than the amount of sterol). This result was previously surmised from semi-quantitative estimates of lipid compositions from area fractions of Lo and Ld phases (43), but the concept that the two phases could be primarily distinguished by their phospholipid content did not gain traction in the community until high quality data from Klaus’ NMR spectrometers were published.

Here, we report a complementary result: demixing of yeast vacuoles into coexisting Lo and Ld phases, which occurs upon a shift from the log to the stationary stage of growth, is accompanied by significant changes in the phospholipid compositions of yeast vacuoles. The fraction of PC lipids roughly doubles. Among the PC lipids, there is a large increase (10°C) in the average *T*_melt_ of the lipids. PE lipids are roughly halved in mole fraction.

In this manuscript, we have analyzed the differences in the vacuole lipidome at two growth stages (the log and the stationary stage) in the context of a single physical variable (the mixing temperature) and how it might affect a single membrane attribute (demixing of the membrane into coexisting liquid phases). Of course, membrane phase separation is only one of many relevant physiological parameters. Lipidomes are also affected by temperature, growth media, carbon source, anoxia, ethanol adaptation, mutations, and cell stress (61, 87, 89–94). Similarly, cells have the potential to adjust their membrane compositions to meet a broad list of inter-related constraints, including lipid packing, thickness, compression, viscosity, permeability, charge, asymmetry, monolayer and bilayer spontaneous curvatures, avoidance of (and proximity to) nonlamellar phases (61, 94–106). Likewise, the lipidome is only one of many biochemical attributes that a cell might vary (e.g. the asymmetry, charge, tension, and adhesion of the membrane; the abundance, crowding, condensation, and crosslinking of proteins; and the conditions of the solvent) to enhance or suppress liquid-liquid phase separation of its membranes (49, 107–111).

Only some of the lipidomic changes that we observe in the vacuole are reflected across the whole cell (18), highlighting the importance of isolating organelles. Our ability to isolate yeast vacuoles with low levels of contamination by non-vacuole organelles leverages recent advances in immunoisolation (25) and our discovery that Mam3 is a robust bait protein in both the log and stationary stages. Recent advances in simulating complex membranes with many lipid types over long time scales are equally exciting (112, 113). We hope that our results will be useful to future modelers in elucidating why stationary stage vacuole membranes phase separate, whereas log stage membranes do not.

## CONCLUSION

Here, we establish that Mam3 is a robust bait protein for immunoisolation of vacuole membranes. Expression levels of Mam3 are high throughout the yeast growth cycle. In the stationary stage, Mam3 partitions into the Ld phase. By conducting lipidomics on isolated vacuole membranes, we find that the shift from the log to the stationary stage of growth is accompanied by large increases in the fraction of PC lipids in the membrane, and that these lipids become longer and more saturated. The resulting increase in melting temperature of these lipids may contribute to demixing of stationary stage vacuole membranes to form coexisting liquid phases.

## SUPPORTING MATERIAL

Supporting material includes a data file, figures, and tables.

## AUTHOR CONTRIBUTIONS

A.J.M., C.E.C., C.L.L., J.R., R.E., and S.L.K. designed research. C.E.C., C.L.L., C.K., and J.R. performed research. C.L.L., J.R., C.K., R.E. and S.L.K. analyzed data. A.J.M., C.L.L., J.R., R.E., and S.L.K. wrote the paper.

## ACKNOWLEDGEMENTS

This research was supported by NSF grant MCB-1925731 to S.L.K, NIH grants GM077349 and GM130644 (to A.J.M.), European Union’s Horizon 2020 research and innovation program (grant agreement no. 866011) to R.E., Deutsche Forschungsgemeinschaft in the framework of the SFB894 to R.E., Volkswagen Foundation grant #93089 to R.E. and J.R., and a travel grant by The Company of Biologists to C.L.L.

This manuscript is dedicated to Klaus Gawrisch, whose contributions to the biophysical community go beyond his rigorous NMR measurements. His open-mindedness to consider and test unpopular ideas, his generosity in sharing credit through collaborations, and his dedication to mentoring of early-career researchers have continuously elevated the quality of scientific results and the collegiality of scientific culture.

## Supporting Material

### Caption for Data File S1

Raw data for abundances of all lipids in vacuoles immuno-isolated with a Mam3 bait protein from yeast in the log stage (left columns) and stationary stage (right columns). At the bottom of the spreadsheet shows sums for each lipid class, and mole percentages. The sums are calculated both with triacylglycerols and ergosterol esters (TAG and EE) and without those storage lipids.

### Supplemental Methods

#### Microscopy

*Saccharomyces cerevisiae* BY4741 (1, 2) expressing a Mam3-GFP fusion protein from its endogenous locus was used for fluorescence microscopy experiments. Live yeast cells were immobilized on a coverslip coated with 3 µL of 1 mg/mL concanavilin-A (EPC Elastin Products Co. catalog no. C2131) in buffer (50 nM Hepes [pH 7.5], 20 mM calcium acetate, and 1 mM MnSO4]. Immediately prior to use, the coverslips were washed with MilliQ water and dried with air. Cells were diluted for imaging in an isosmotic solution of conditioned medium from the grown culture. To produce conditioned medium, 1 mL of culture was centrifuged at 3,400 × g. The supernatant was collected and centrifuged again at 3,400 × g to remove remaining cells. To decrease refractive index mismatch (3), 200 µL of OptiPrep (60% OptiPrep Density Gradient Medium; Sigma catalog no. D1556) was added to 800 µL of medium and vortexed. Samples of 3 µL of cells were diluted into 3 µL of conditioned medium containing 12% OptiPrep and placed onto the concanavalin-A–coated coverslip. Cells were sandwiched with a top coverslip and allowed to adhere to the coated coverslip for 10 min before imaging. Unless otherwise noted, images were acquired on a Nikon TE2000 microscope equipped with a Teledyne Photometrics Prime 95BSI camera. Using an oil-immersion objective (100×, 1.4 numerical aperture), GFP was excited with X-Cite 110 light-emitting diode light source and filtered through an infrared cut filter to prevent aberrant heating of the sample from the optics. Brightness and contrast were adjusted linearly using ImageJ software (https://imagej.nih.gov/ij/).

#### Preparation of magnetic beads

Dynabeads with Protein G (Thermo Fisher Scientific #10009D) from 1.6 mL of slurry were washed with 1.6 mL PBS-T. After resuspension in 1.6 mL fresh PBS-T, 10 µL of anti-FLAG antibody (M2, monoclonal mouse IgG1, affinity isolated, F1804, 1 mg/ml) was added to the Dynabeads. This results in a sub-saturated coverage of Dynabeads with antibody. The resulting mix was incubated overnight at 4°C and 20 rpm overhead rotation. The supernatant was removed and the antibody-coated magnetic beads were washed once with 1.6 mL of PBS-T and twice with 1.6 mL IP buffer (25 mM HEPES pH 7.0, 1 mM EDTA, 150 mM NaCl) before use in immuno-isolation.

#### Lipid extraction, lipidomics data acquisition and post-processing

Mass spectrometry-based shotgun lipidomics was performed by Lipotype GmbH (Dresden, Germany) as described (4, 5). Lipids were extracted using a two-step chloroform/methanol procedure (4). Samples were spiked with internal lipid standards for major lipid classes in which the lipids contain combinations of acyl chains not found in biological samples, as previously described (6). After extraction, the organic phase was transferred to an infusion plate and dried in a speed vacuum concentrator. 1st step dry extract was re-suspended in 7.5 mM ammonium acetate in chloroform/methanol/propanol (1:2:4, V:V:V) and 2nd step dry extract in 33 % ethanol solution of methylamine in chloroform/methanol (0.003:5:1; V:V:V). All liquid handling steps were performed using Hamilton Robotics STARlet robotic platform with the Anti Droplet Control feature for organic solvents pipetting.

Samples were analyzed by direct infusion on a QExactive mass spectrometer (Thermo Scientific) equipped with a TriVersa NanoMate ion source (Advion Biosciences). Samples were analyzed in both positive and negative ion modes with a resolution of Rm/z=200=280000 for MS and Rm/z=200=17500 for MSMS experiments, in a single acquisition. MS/MS was triggered by an inclusion list encompassing corresponding MS mass ranges scanned in 1 Da increments (7). Both MS and MSMS data were combined to monitor ergosterol-esters (EE), diacylglycerol (DAG) and triacylglycerol (TAG) ions as ammonium adducts; Phosphatidylcholine (PC) as an acetate adduct; and cardiolipin (CL), phosphatidic acid (PA), phosphatidylethanolamine (PE), phosphatidylglycerol (PG), phosphatidylinositol (PI) and phosphatidylserine (PS) as deprotonated anions. MS only was used to monitor the lyso-lipids of PA, PE, PI, and PS, as well as inositolphosphorylceramide (IPC), mannosyl-inositolphosphorylceramide (MIPC), and mannosyl-di-(inositolphosphoryl)ceramide (M(IP)2C) as deprotonated anions; ceramine (Cer) and lyso-PC (LPC) as acetate adducts and ergosterol as protonated ion of an acetylated derivative (8).

Data were analyzed with in-house developed lipid identification software based on LipidXplorer (9, 10). Data post-processing and normalization were performed using an in-house developed data management system. Only lipid identifications with a signal-to-noise ratio >5, and a signal intensity 5-fold higher than in corresponding blank samples were considered for further data analysis.

#### Microsomal preparation

Throughout microsomal preparation and immuno-isolation procedures (Fig. 2 of the main text), samples from supernatants and pellets were retained for immunoblot analysis. Break points are indicated where samples can be stored at −80°C.

4,000 OD_600_·mL of yeast cells were required to yield a sufficient biomass as starting material for isolating vacuole membranes. Cells were harvested by centrifugation at 3,000 x g for 5 min at room temperature. The resulting pellet was resuspended in 25 mL of pre-cooled phosphate buffered saline (PBS) and placed on ice. Cell suspensions corresponding to 1,000 OD_600_ mL were transferred to 50 ml tubes and sedimented at 3,000 x g for 5 min at 4°C. The supernatants were discarded, and the cell pellets were snap frozen in liquid nitrogen and stored at −80°C.

A cell pellet of 1,000 OD_600_·mL was thawed from −80°C slowly on ice and resuspended in 10 mL of microsome preparation (MP) buffer (25 mM HEPES at pH 7.0, 1 mM EDTA, 0.6 M mannitol, to which 30 µg/ml protease inhibitor cocktail (10 µg/mL of pepstatin, antipain, chymotrypsin each) and 12.5 units/mL benzonase was added freshly). To mechanically break the cells (Fig. 2A), 13 g pre-chilled zirconia glass beads (0.5 mm diameter) were combined with the resuspended cell pellet in a 15 mL tube. The tube was filled to the top with MP buffer to displace all remaining air that would give rise to air bubbles during mechanical agitation. Using a FastPrep-24 bead beater at 4°C, cells were subjected to 10 cycles of shaking for 15 s at 5 m/s, followed by 45 s of cooling on ice. The resulting cell lysates were transferred to fresh 15 mL tubes. 2 mL MP buffer were used to wash the zirconia glass beads and then combined with the previous lysates.

Differential centrifugation was performed to separate a crude microsomal membrane fraction from cell debris and the soluble proteins leading to an enrichment of vacuole membranes (Fig. 2A). Cell lysates were spun at 3,234 x g and 4°C for 5 min in a swinging bucket rotor. The supernatant was transferred to a fresh 15 ml tube and re-centrifuged at 3,234 x g and 4°C for 5 min. The resulting supernatant (S3) was then transferred to ultracentrifuge bottles (26.3 mL polycarbonate bottle assemblies, Beckman Coulter #355618), balanced with MP buffer and centrifuged (rotor Type 70 Ti) at 12,000 x g and 4°C for 20 min. The resulting supernatant (S12) was transferred to a fresh ultracentrifuge bottle, balanced with MP buffer, and centrifuged at 100,000 x g at 4°C for 60 min. Although vacuole markers Vph1 and Vac8 are also found in the pellet after 12,000 x g centrifugation (P12) and in the supernatant after 100,000 x g centrifugation (S100), we chose to work with the pellet after 100,000 x g centrifugation (P100) because it contains smaller vesicles that are less likely connected to mitochondria and lipid droplets. To avoid contamination of the microsome pellet (P100) by lipid droplets floating on top of the supernatant (S100), the supernatant was removed by vacuum from the ultracentrifuge tube, working carefully from the top to the bottom. The pellet containing crude microsomes (P100) was rinsed with 15 mL of MP buffer to remove remnants of the supernatant before resuspending it in 1 mL MP buffer, snap freezing in liquid nitrogen, and storage at −80°C.

Pellets from the microsomal preparation were thawed from −80°C slowly on ice and then sonicated to segregate aggregated membrane vesicles and to break vacuoles into smaller vesicles (Fig. 2A of the main text) (6). A tip sonicator (MS72 sonotrode on a Bandelin Sonopuls HD 2070) was used for 10 s, with a duty cycle of 0.7 at 50% amplitude, keeping the samples on ice. Previous experiments revealed that this treatment does not cause membrane mixing (6). Sonication transformed the solution from cloudy to clear. The solution was then centrifuged at 3,000 x g at 4°C for 3 min. The supernatant, which contained vesicles, was used for subsequent immuno-isolation and the pellet was discarded.

---

**Figure S1:**
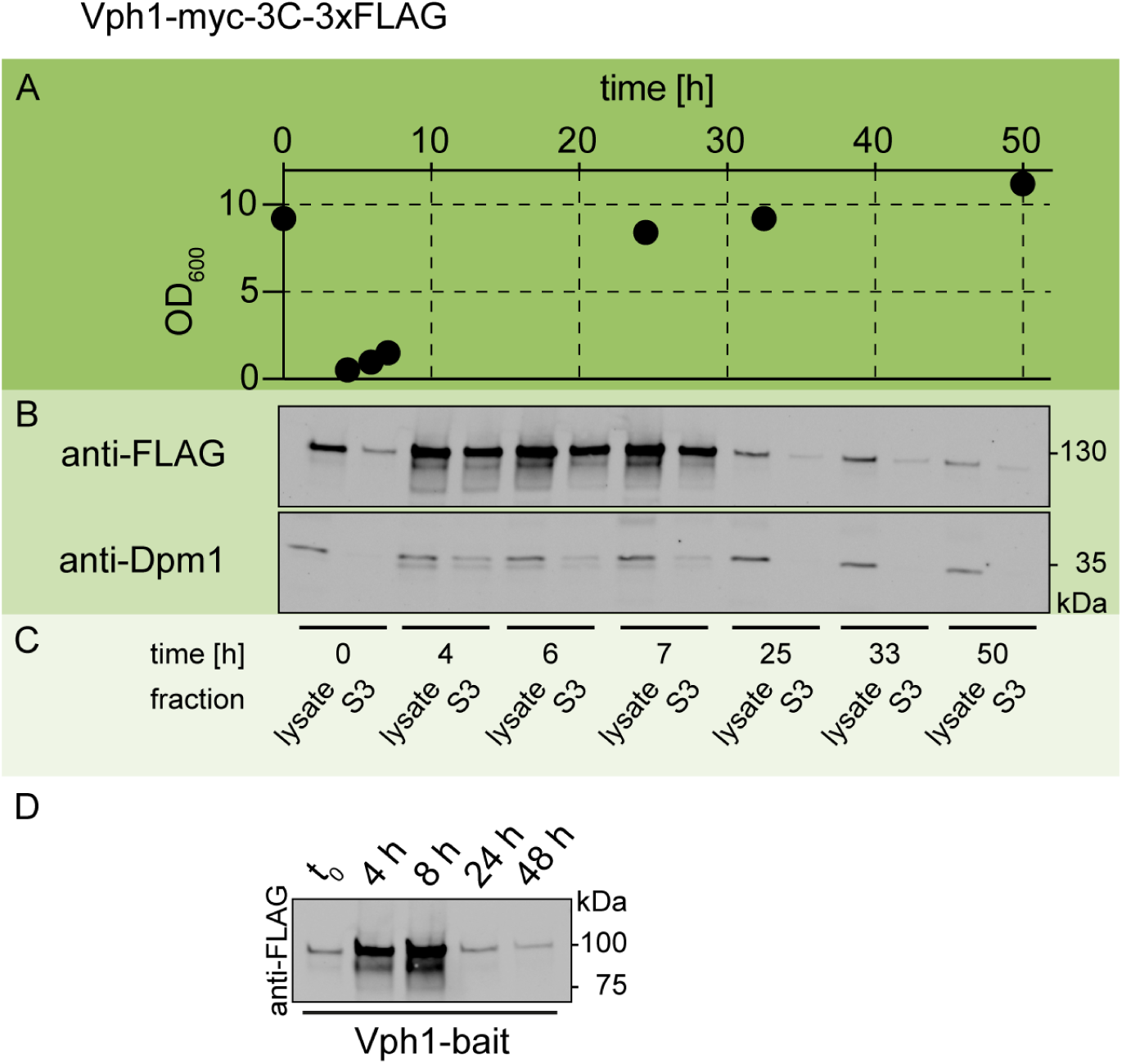
Expression levels of the Vph1-bait construct (Vph1-myc-3C-3xFLAG) are low in the stationary stage. **(A)** Yeast growth curve through time, where OD_600_ is the optical density at 600 nm, a proxy for the density of cells. The sample at time “0” is a preculture after 19 hours of the preculture’s growth. The preculture was then diluted to OD_600_ = 0.1 and allowed to grow into the log (early time points), and stationary stage (late time points). **(B-C)** At each timepoint, an immunoblot was performed for the cell lysate (left columns) and supernatant (“S3”, right columns). The supernatant corresponds to S3 in Fig. 2, which is collected from the supernatant after the first step of microsome preparation, a 3,000 x g centrifugation. The presence of Vph1 was visualized by staining for an FLAG tag attached to it. In the log stage, Vph1 is abundant (timepoints 1-3) in the cell lysate and S3 samples. This result is consistent with reported values of > 50,000 copies/cell, which likely applies only to the log stage (up to ∼10 hours of growth) (11, 12). In the stationary stage (timepoints 4-6), Vph1 is significantly reduced in the cell lysate and further reduced after centrifugation step S3. Dpm1, a protein that is a marker for the endoplasmic reticulum, is used as a control. Dpm1 is reported to have < 2,000 copies/cell (11). It maintains a roughly constant abundance throughout the growth cycle. **(D)** A minimal experiment reproducing the results in panel B, with explicit values for time points and molar masses.

**Figure S2:**
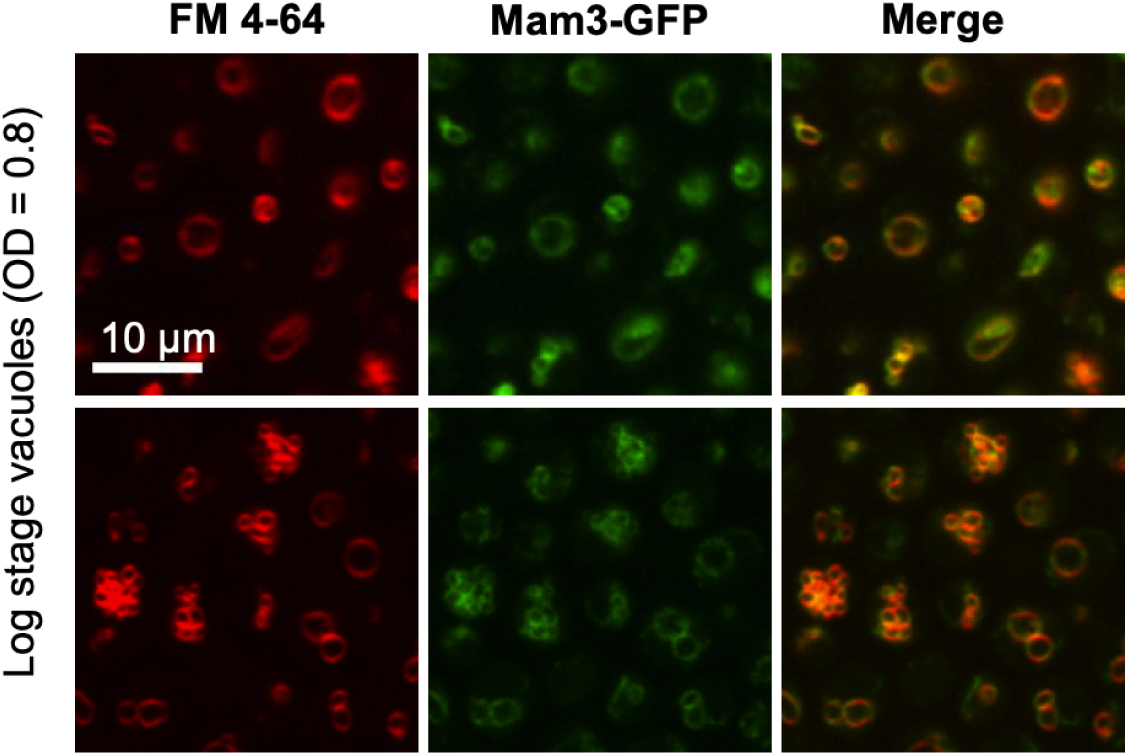
The protein Mam3-GFP (middle column) colocalizes with FM 4-64 (left column), which is known to label vacuole membranes. All yeast in the field of view are living and are in the logarithmic stage; no micron-scale domains appear in their membranes. Images were taken at room temperature.

**Figure S3:**
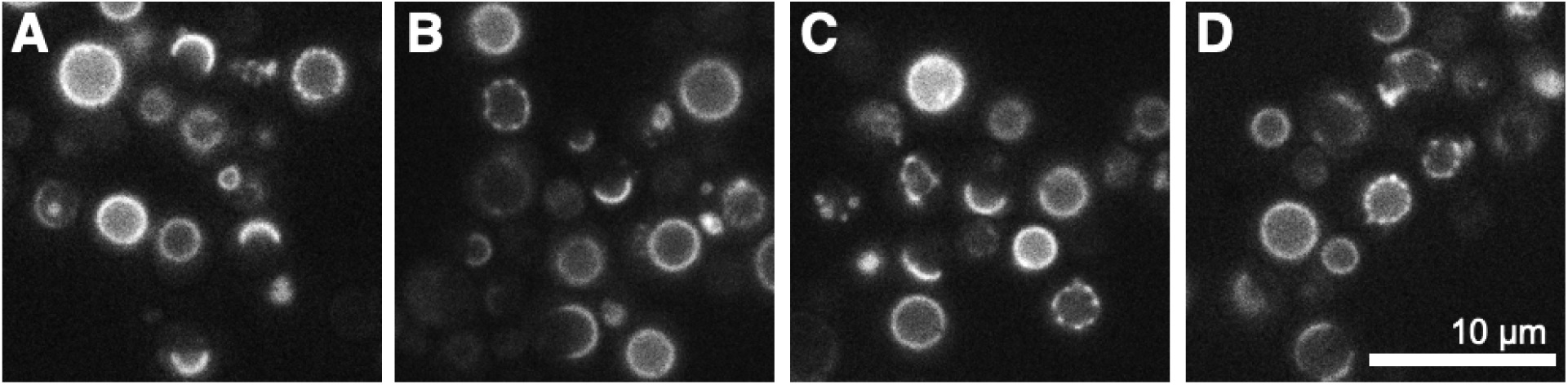
Four fields of view of vacuole membranes in living yeast cells in the stationary stage after 48 hours of cultivation, under equivalent conditions. Bright areas of the membranes contain the fluorescent protein Mam3-GFP, which was endogenously labeled. Most membranes in the fields of view have phase-separated into micron-scale domains. Images were taken at room temperature.

**Figure S4:**
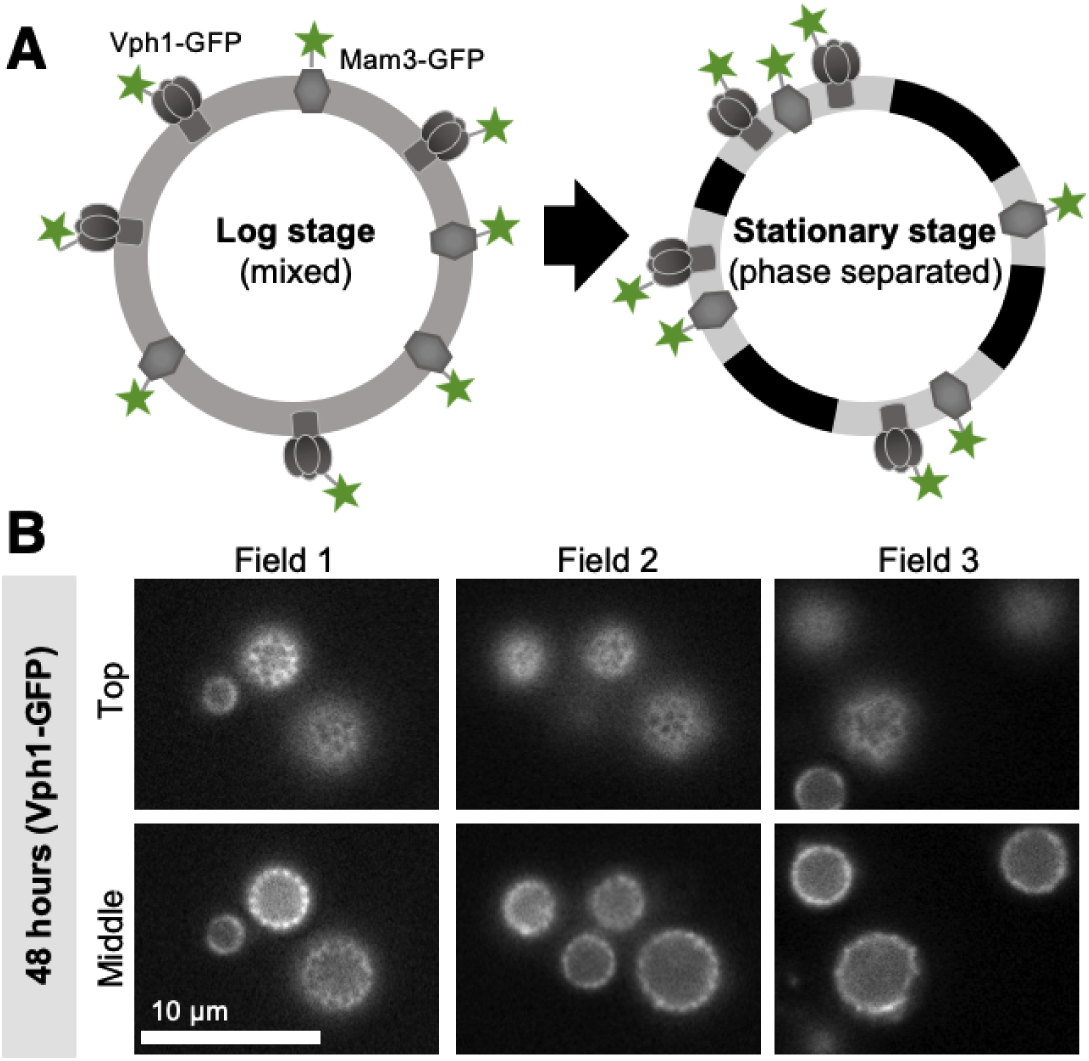
**(A) Left:** For yeast in the logarithmic stage of growth, lipids and proteins appear uniformly distributed across the surface of vacuole membranes. Two of the proteins, Vph1 and Mam3, can be endogenously labeled with GFP (as shown here) or a bait tag for immunoisolation. **Right:** In the stationary stage, the membrane separates into two liquid phases. Vph1-GFP and Mam3-GFP preferentially partition to the same phase. **(B)** After 48 hours of growth, yeast are in the stationary stage of growth and most vacuole membranes have phase separated. The proteins Mam3 (Fig. 3) and Vph1 partition into only one of the phases, shown at both the top and the midplane of the vacuoles in each field of view. Images are of living cells and were taken at room temperature.

**Figure S5:**
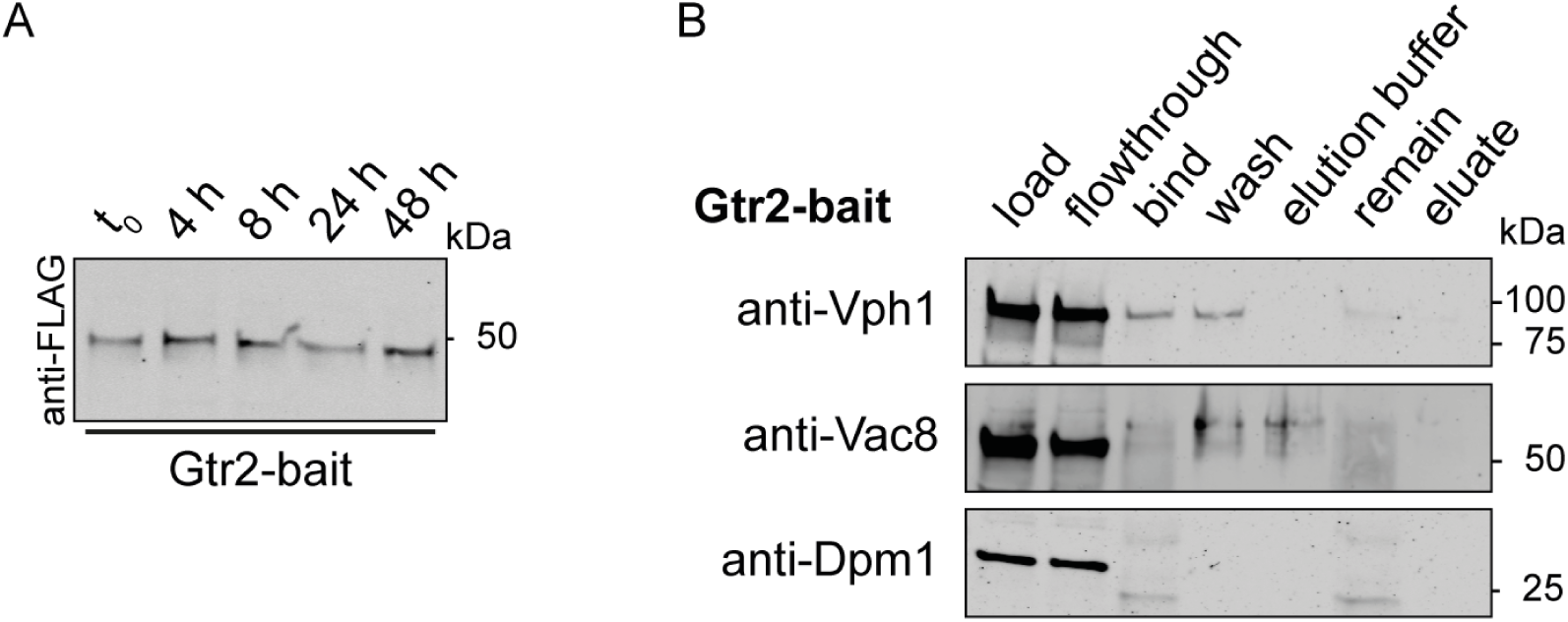
Gtr2 is a membrane associated protein anchored to the vacuole membrane via a lipid anchor. In phase separated stationary stage vacuoles, it resides in the Lo phase (13). **(A)** The Gtr2-bait protein is equally abundant throughout the logarithmic and stationary growth stages. **(B)** We were not able to immuno-isolate any membranes via the Gtr2-bait protein performing MemPrep as described in Fig. 2. In immunoblot analysis, the vacuole markers Vph1 and Vac8 are not detectable in the eluate fraction. A corresponding figure for Mam3 protein is in Fig. 4 of the main text.

**Figure S6:**
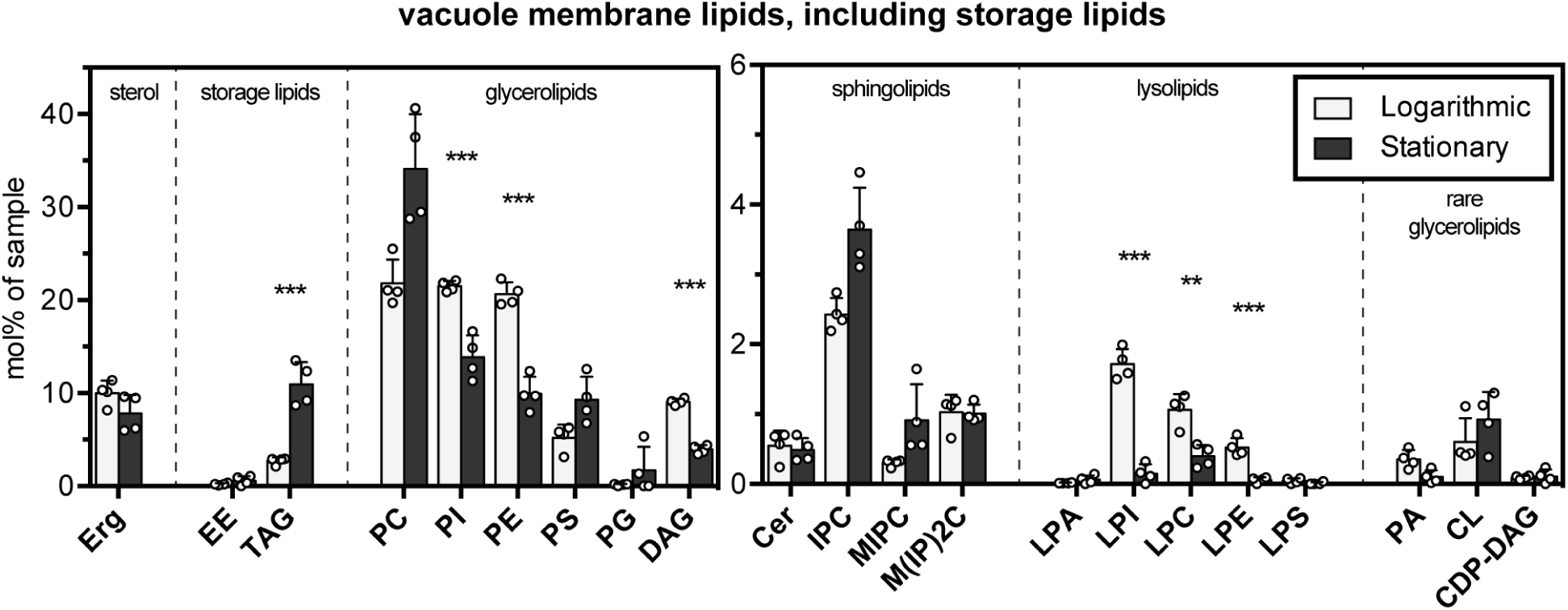
Lipids in log stage and stationary stage vacuole membranes immuno-isolated with a Mam3 bait tag; note that the y-axis range is smaller in the graph at the right. Acronyms of lipid names are listed in Table S1. Error bars are standard deviations of vacuole samples immuno-isolated on different days. Data in this figure include the storage lipids of ergosterol esters (EE) and triacylglycerols (TAG), which are predominantly found in lipid droplets. For yeast in the log stage of growth, we find that ergosterol esters constitute 0.3% of all isolated phospholipids, ergosterol, and ergosterol esters, in contrast to 4.2% found by Zinser et al. (14). Specifically, Zinser et al. find a mole ratio of ergosterol to phospholipids of 0.18 and a mole ratio of ergosterol ester to ergosterol of 0.29 (14). Statistical significance was tested by multiple t-tests correcting for multiple comparisons (method of Benjamini et al. (15)), with a false discovery rate Q = 1%, without assuming consistent standard deviations. *p < 0.05, **p < 0.01, ***p < 0.001.

**Figure S7:**
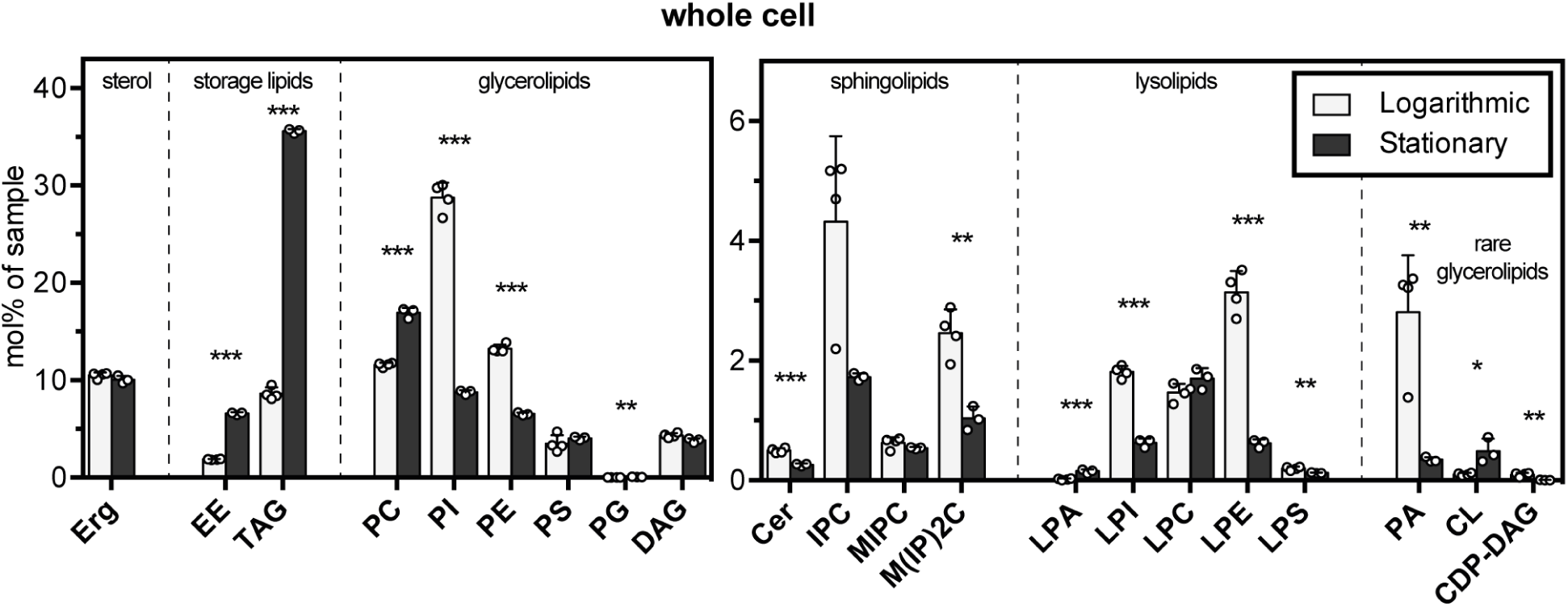
Lipidome of whole yeast cells in the logarithmic and the stationary stage. Error bars represent standard deviations of four independent experiments in the logarithmic stage and three independent experiments in the stationary stage. The data from logarithmic cells are taken from (6). Statistical significance was tested by multiple t-tests correcting for multiple comparisons (method of Benjamini et al. (15)), with a false discovery rate Q = 1%, without assuming consistent standard deviations. *p < 0.05, **p < 0.01, ***p < 0.001.

**Figure S8:**
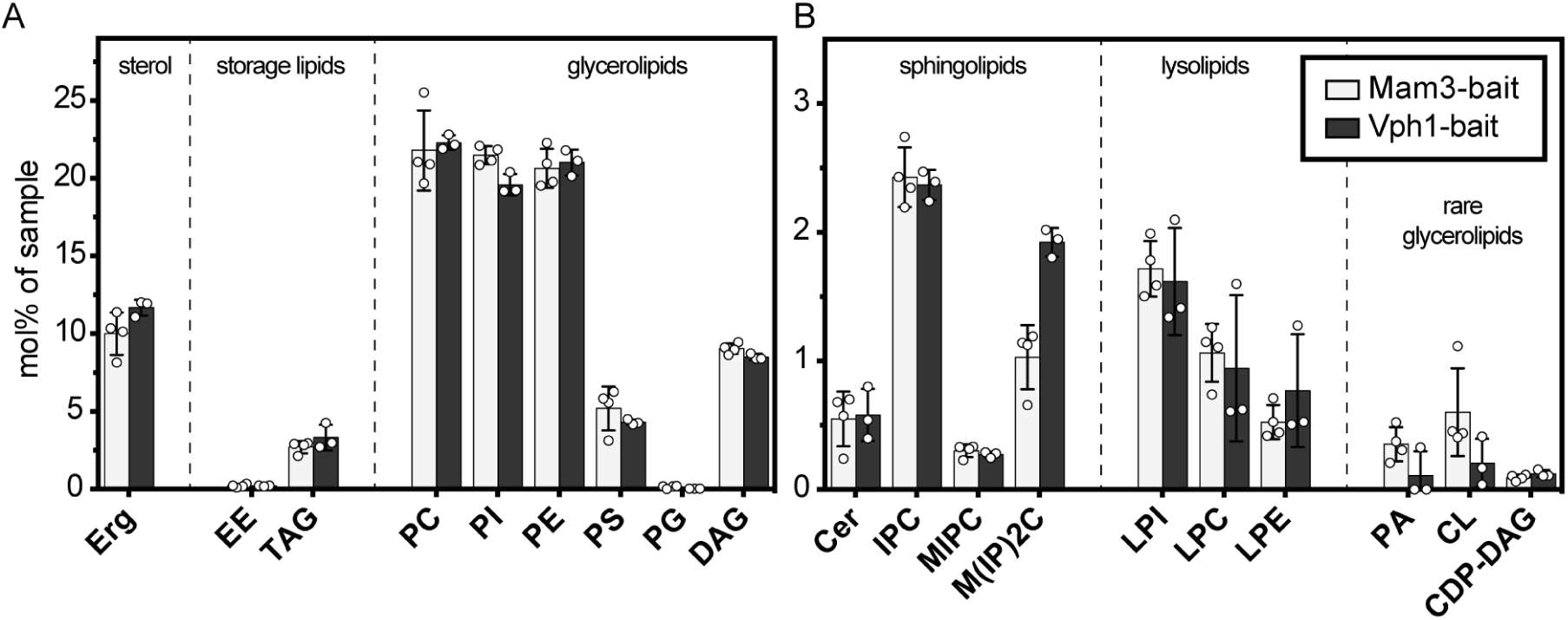
Lipidome of vacuole membranes from yeast in the log stage isolated by two different protein bait tags: Mam3 and Vph1. Vph1 data is from (6). Error bars are standard deviations of vacuole samples immuno-isolated on different days. Overall, the two lipidomes are in close agreement. The only apparent differences are in the amount of M(IP)2C (which may be inconsequential because it is in low abundance and which can change rapidly in its abundance during the log stage of growth (16)) and in PS levels. These differences might reflect real disparities in how vacuole membranes are isolated by Vph1 and Mam3 baits, or they might reflect experimental variation. Full Vph1 data are presented and discussed in (6). **(A)** Three classes of lipids (glycerolipids, sterol and storage lipids) represent a majority of the lipidome. **(B)** Other classes of lipids (e.g. sphingolipids and ceramides) are much less abundant. Note that the y-axis range is roughly an order of magnitude smaller in panel B than in panel A. Statistical significance was tested by multiple t-tests correcting for multiple comparisons using the method of Benjamini et al. (15), with a false discovery rate Q = 1%, without assuming consistent standard deviations. All differences are non-significant.

**Figure S9:**
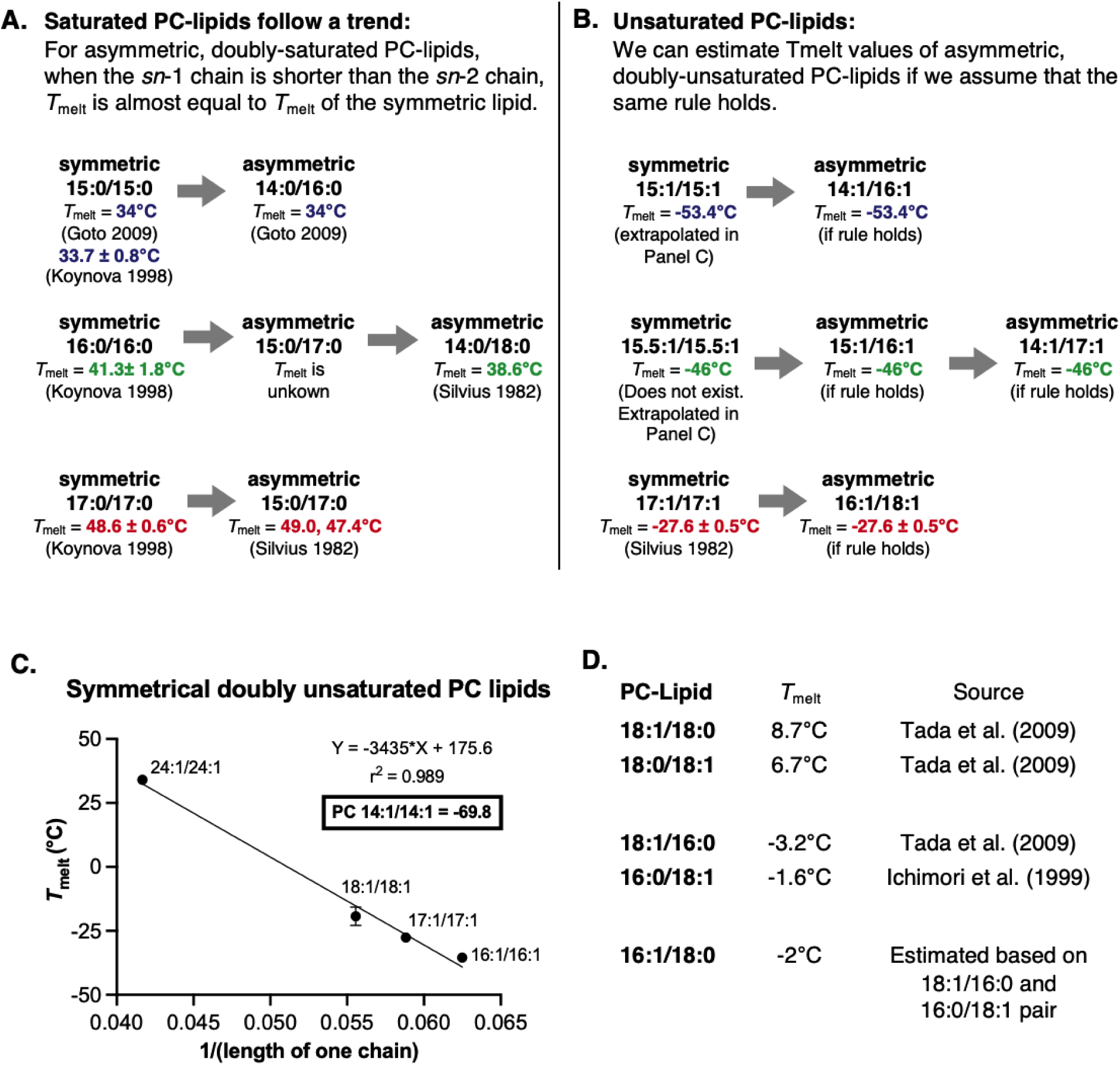
Experimental trends used to estimate melting temperatures for PC-lipids.

**Figure S10:**
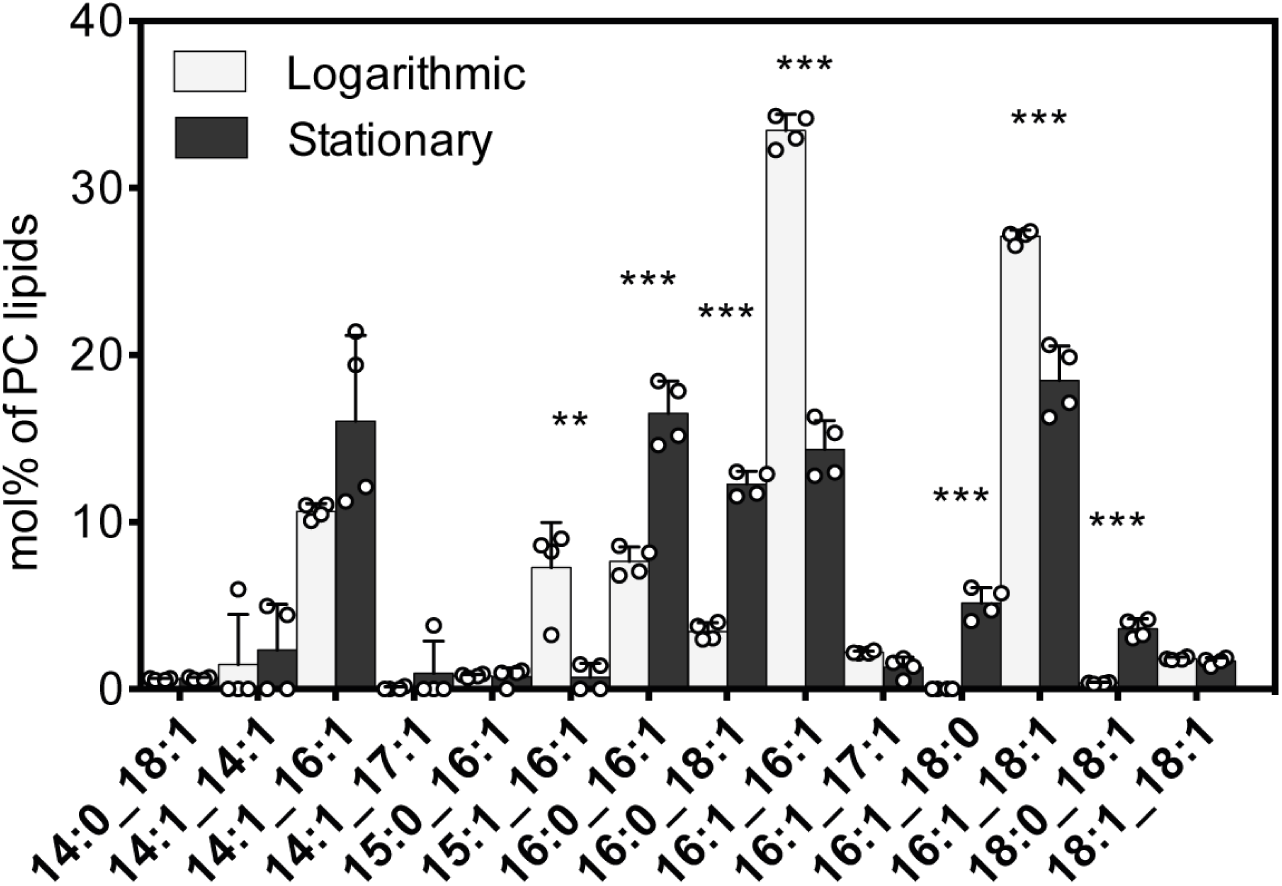
Mole percent of PC lipids with different acyl chains in immuno-isolated vacuole membranes from yeast in the logarithmic and stationary stages. The four data points superimposed on each bar are from four independent experiments. Statistical significance was tested by multiple t-tests correcting for multiple comparisons (method of Benjamini et al. (15)), with a false discovery rate Q = 1%, without assuming consistent standard deviations. *p < 0.05, **p < 0.01, ***p < 0.001.

**Figure S11:**
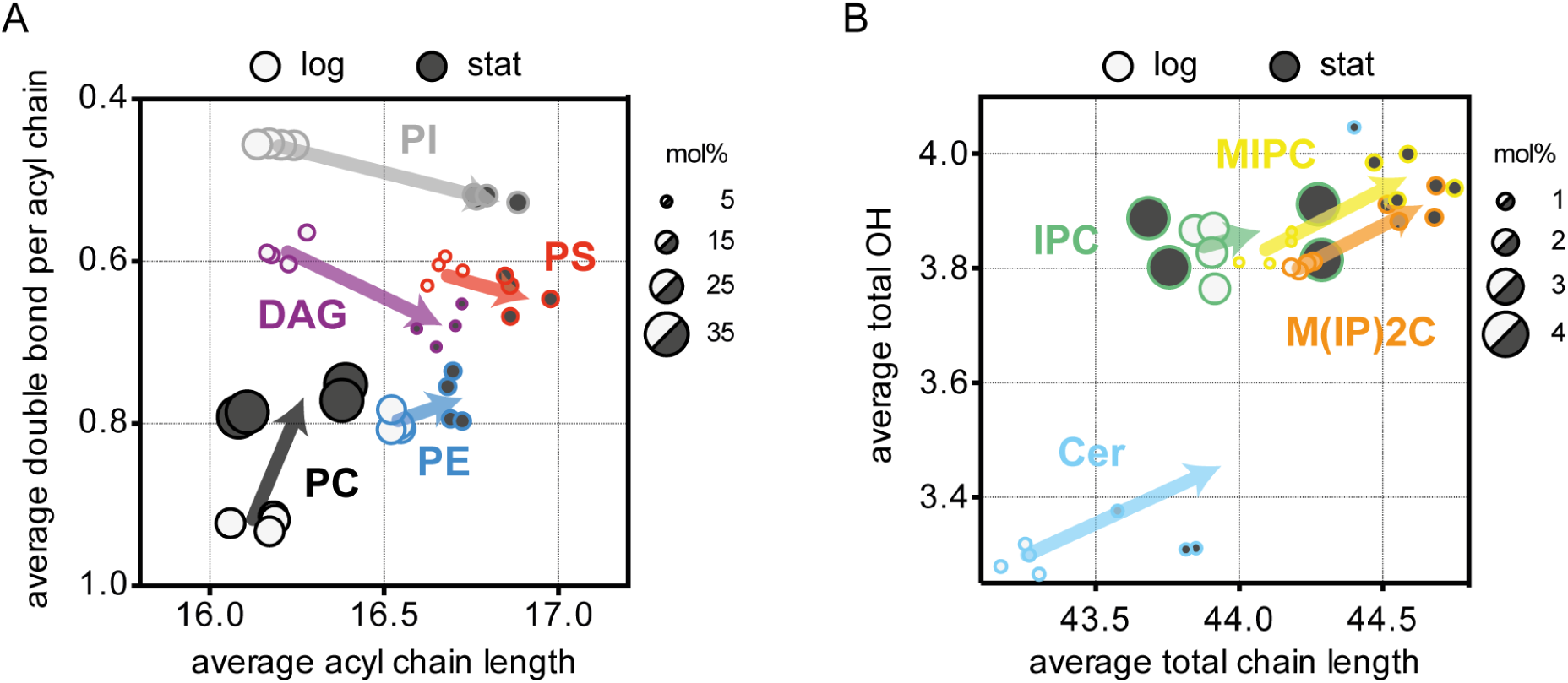
(A) Average number of double bonds per acyl chain and average acyl chain length in the log (open circle) and stationary stage (filled circle) for several lipid types. The size of the circles represents the mole percent of those lipids in immuno-isolated vacuole membranes, excluding storage lipids such as triacylglycerols and ergosterol esters. From bottom to top, data for PC, PE, DAG, PS, and PI lipids are shown. Four data sets are shown, from four independent experiments. The arrows show the average change from the log to the stationary stage. (B) The same type of graph as in panel A is repeated for Cer, IPC, MIPC, and M(IP)_2_C lipids, except that the y-axis shows the average total number of OH groups for the entire lipid and the x-axis shows the total chain lengths of respective sphingolipids.

**Figure S12:**
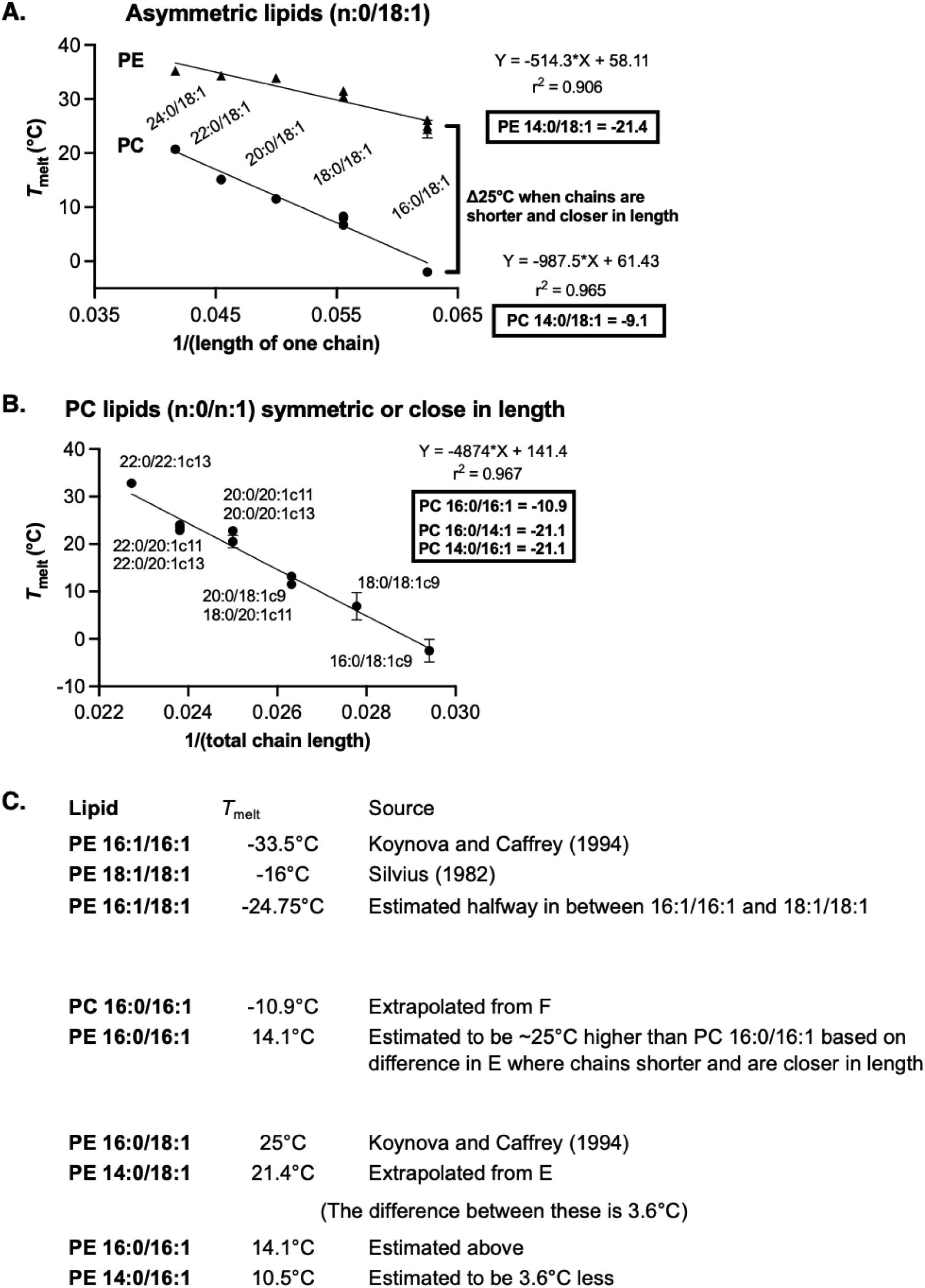
Trends used to estimate melting temperatures for PE-lipids.

**Figure S13:**
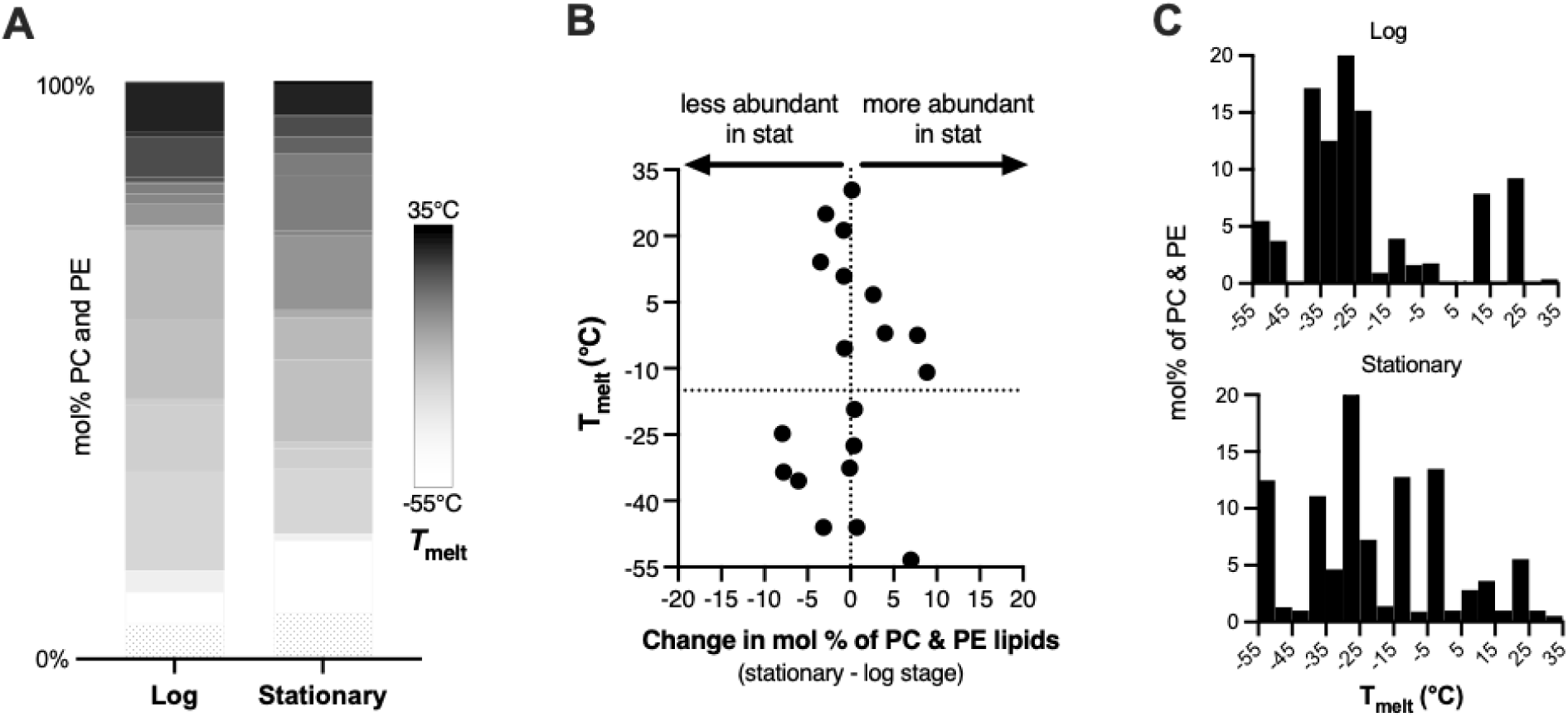
(A) Distributions of melting temperatures for PC and PE lipids of yeast vacuoles, in ratios corresponding to log and stationary stage vacuoles. In other words, the data are weighted to account for the increase in PC lipids and the decrease in PE lipids from the log stage to the stationary stage. (B) Changes in mol% of PC and PE lipids (x-axis) for each melting temperature (y-axis). The change in average lipid *T*_melt_ due to PC lipids is offset by a decrease in the fraction of PE lipids. (C) Histograms of the percent of PC and PE lipids at each melting temperature, in the log and stationary stages of growth. If we treat the data in the histograms as if they are independent points in a scatterplot, then the values would have standard deviations of 17°C in the log stage and 21°C in the stationary stage.

**Table S1:** Mole percent phospholipids in log-stage, isolated vacuole membranes. (Renormalized to sum all phospholipids to 100%)

| **Source** | **PC** | **PE** | **PI** | **PS** | **PA** | **Lyso** | **CL** | **Other** | **Method** |
| --- | --- | --- | --- | --- | --- | --- | --- | --- | --- |
| Zinser 1991 | 46.5 | 19.4 | 18.3 | 4.4 | 2.1 |  | 1.6 | 7.7 | *density gradient* |
|  | 39.2 | 26.6 | 24.4 | 3.9 | 2.5 | 1.6 | 0.4 | 1.4 |  |
| Tuller 1999 | $\pm 3.3$ | $\pm 1.5$ | $\pm 1.8$ | $\pm 1.0$ | $\pm 0.8$ | $\pm 0.8$ | $\pm 0.1$ | $\pm 0.1$ | *density gradient* |
|  | 31.4 | 29.6 | 27.6 | 6.1 | 0.2 | 4.7 | 0.3 | 0.2 |  |
| Reinhard 2022 | $\pm 1.0$ | $\pm 1.5$ | $\pm 0.6$ | $\pm 0.3$ | $\pm 0.3$ | $\pm 1.9$ | $\pm 0.3$ | $\pm 0.0$ | *immuno-isolation* |
|  | 29.6 | 28.0 | 29.2 | 7.1 | 0.5 | 4.5 | 0.8 | 0.3 |  |
| This work | $\pm 2.7$ | $\pm 1.0$ | $\pm 1.4$ | $\pm 2.1$ | $\pm 0.2$ | $\pm 0.2$ | $\pm 0.4$ | $\pm 0.1$ | *immuno-isolation* |

**Conditions for the experiments above:**
|  | **Strain** | **Glucose** | **Temp** | **Media** | **OD** | **Stage/Hours** |
| --- | --- | --- | --- | --- | --- | --- |
| Zinser 1991 | *X-2180* | 3% | — | YPD | — | — |
| Tuller 1999 | *FY1679* | 2% | 30°C | YPD | ? | *Late log (~16 h)* |
| Reinhard 2022 | *BY4741* | 2% | 30°C | SCD | 1 | *Mid log (8 h)* |
| This work | *BY4741* | 2% | 30°C | SCD | 1 | *Mid log (8 h)* |

**Notes:** Uncertainties are standard deviations. Zinser et al. did not provide values of uncertainties (14). Tuller et al. note that FY1679 cells are atypical in their high fractions of palmitic acid (16:0) carbon chains and low fraction of oleic acid (18:1) chains (17), implying that comparisons between strains may not always be valid. A contamination of 0.4% cardiolipin in vacuoles isolated by Tuller et al. represents 5.5% contamination by mitochondrial membranes (of which $7.2 \pm 0.2\%$ of lipids are cardiolipin) (17). In the row titled “This work”, the entry for “Other” denotes PG lipids. If sphingolipids (IPC, MIPC, and MI(IP)2C) are included in “Other”, the mole percent of other lipids for this work increases to $5.0 \pm 0.8\%$

**Table S2:** Values of T_melt_ for lipids in yeast vacuole membranes.

| **PC Lipid:** | **$T_{\text{melt}}$ (°C)** | **Source of $T_{\text{melt}}$ value** |
| --- | --- | --- |
| **14:1/14:1** | -69.8 | Extrapolated in Fig. S9, panel C |
| **14:1/16:1** | -53.4 | Estimated in Fig. S9, panel B |
| **14:1/17:1** | -53.4 | Estimated in Fig. S9, panel B |
| **15:1/16:1** | -46.0 | Estimated in Fig. S9, panel B |
| **16:0/16:1** | -10.9 | Estimated in Fig. S9, panel B |
| **16:0/18:1** | -2.5 ± 2.4 | Koynova & Caffrey (1998), (18) |
| **16:1/16:1** | -35.5 | Silvius (1982), (19) |
| **16:1/17:1** | -32.6 | Estimated using the logic of Fig. S9, panel B |
| **16:1/18:0** | -2 | Estimated in Fig. S9, panel D |
| **16:1/18:1** | -27.5 ± 0.6 | Silvius (1982), (19) |
| **18:0/18:1** | 6.7 | Tada (2009), (20) |
| **18:1/18:1** | -19.3 ± 3.6 | Silvius (1982),(19) |

| **PE Lipid:** | **$T_{\text{melt}}$ (°C)** | **Source of $T_{\text{melt}}$ value** |
| --- | --- | --- |
| **14:0/16:1** | 10.5 | Estimated in Fig. S11, panel C |
| **14:0/18:1** | 21.4 | Extrapolated in Fig. S11, panel A |
| **16:0/16:1** | 14.1 | Estimated in Fig. S11, panel C |
| **16:0/18:1** | 25 | Wang (1994), (21) |
| **16:1/16:1** | -33.5 | Koynova & Caffrey (1994), (22) |
| **16:1/18:1** | 19.5 | Estimated in Fig. S11, panel C |
| **18:0/18:1** | 30.4 | Silvius (1982), (19) |
| **18:1/18:1** | -5.5 ± 0.5 | Matsuki et al. (2017), (23) |

**Table S3:** Literature values of *T*_melt_ for PC-lipids. (Used for estimating *T*_melt_ values for vacuole lipids)

| Chain length : unsaturation | $T_{\text{melt}}$ (°C) | Reference | Reference # |
| --- | --- | --- | --- |
| 13:0/13:0 PC | 13.5 | Silvius (1982) | (19) |
| 14:0/14:0 PC | 24 | Goto et al. (2009) | (24) |
| 15:0/15:0 PC | 34<br>33.7 ± 0.8 | Goto et al. (2009)<br>Koynova & Caffrey (1998) | (18, 24) |
| 16:0/16:0 PC | 41.3 ± 1.8 | Koynova & Caffrey (1998) | (18) |
| 17:0/17:0 PC | 48.6 ± 0.6 | Koynova & Caffrey (1998) | (18) |
| 14:0/16:0 PC | 34 | Goto et al. (2009) | (24) |
| 14:0/18:0 PC | 38.6 | Silvius (1982) | (19) |
| 15:0/17:0 PC | 49.0, 47.4 | Silvius (1982) | (19) |
| 16:1/16:1 PC | -35.5, -36 | Silvius (1982) | (19) |
| 17:1/17:1 PC | -27.6 ± 0.5 | Silvius (1982) | (19) |
| 18:1/18:1 PC | -19.3 ± 3.6 | Silvius (1982) | (19) |
| 24:1c9/24:1c9 PC | 34 | Koynova & Caffrey (1998) | (18) |
| 16:0/18:1 PC | -1.6 | Ichimori et al. (1999) | (25) |
| 18:1/16:0 PC | -3.2 | Tada et al. (2009) | (20) |
| 18:0/18:1 PC | 6.7 | Tada et al. (2009) | (20) |
| 18:1/18:0 PC | 8.7 | Tada et al. (2009) | (20) |
| 20:0/18:1 PC | 11.5 ± 0.5 | Koynova & Caffrey (1998) | (18) |
| 22:0/18:1 PC | 15.1 | Koynova & Caffrey (1998) | (18) |
| 18:0/20:1c11 PC | 13.2 | Koynova & Caffrey (1998) | (18) |
| 20:0/20:1c11 PC | 20.5 ± 1.3 | Koynova & Caffrey (1998) | (18) |
| 20:0/20:1c13 PC | 22.8 | Koynova & Caffrey (1998) | (18) |
| 22:0/20:1c11 PC | 22.9 | Koynova & Caffrey (1998) | (18) |
| 22:0/20:1c13 PC | 23.5, 24 | Koynova & Caffrey (1998) | (18) |
| 22:0/22:1c13 PC | 32.8 | Koynova & Caffrey (1998) | (18) |

**Table S4:** Literature values of T_melt_ for PE-lipids. (Used for estimating *T*_melt_ values for vacuole lipids)

| Chain length : unsaturation | $T_{\text{melt}}$ (°C) | Reference | Reference # |
| --- | --- | --- | --- |
| 16:0/18:1 PE | 24.41 ± 1.63<br>26.1 | Koynova & Caffrey (1994)<br>Wang et al. (1994) | (18) |
| 18:0/18:1 PE | 31.5<br>30.4 | Wang et al. (1994)<br>Silvius (1982) | (19, 21) |
| 20:0/18:1 PE | 33.9 | Wang et al. (1994) | (21) |
| 22:0/18:1 PE | 34.3 | Wang et al. (1994) | (21) |
| 24:0/18:1 PE | 35.2 | Wang et al. (1994) | (21) |
| 16:1/16:1 PE | -33.5 | Koynova & Caffrey (1994)<br>Silvius (1982) | (18)<br>(19) |
| 18.1/18.1 PE | -5.5 ± 0.5 | Matsuki et al. (2017) | (23) |

**Table S5:** How changes in lipid chain length and unsaturation affect *T*_melt_. (Values are from Tables S2-S4)

| Headgroup | Lipid 1, $T_{\text{melt}}$ | Lipid 2, $T_{\text{melt}}$ | Change in the<br>average number<br>of carbons<br>(over both chains) | Change in<br>the average<br>saturation<br>(over both chains) | Change<br>in<br>$T_{\text{melt}}$<br>(per lipid) |
| --- | --- | --- | --- | --- | --- |
| PC | di(16:0)PC, 41.3°C | di(17:0)PC, 48.6°C | 2 | – | 7.3°C |
| PC | 16:0/18:1PC, -1.6°C | 18:0/18:1PC, 6.7°C | 2 | – | 8.3°C |
| PE | 16:0/18:1PE, ~25°C | 18:0/18:1PE, ~31°C | 2 | – | 6°C |
| **AVERAGE EFFECT OF ADDING ONE CARBON PER LIPID** |  |  |  |  | **~ 7.2°C** |
| PC | 18:1/18:1PC, -19.3°C | 18:0/18:1PC, 6.7°C | – | 1 | 26°C |
| PC | 18:1/18:1PC, -19.3°C | 18:1/18:0PC, 8.7°C | – | 1 | 28°C |
| PE | 18:1/18:1PE, -5.5 | 18:0/18:1PE, 31.0 ± 0.8°C | – | 1 | 37°C |
| **AVERAGE EFFECT OF ADDING ONE SATURATED BOND PER LIPID** |  |  |  |  | **~ 30.3°C** |

**Table S6:** Differences in length of sn-1 and sn-2 chains of lipids from vacuoles in the log and stationary stages.

| **Difference in number of carbons** | **Log stage (average ± standard deviation)** | **Stationary stage (average ± standard deviation)** |
| --- | --- | --- |
| 0 carbons | 41.0 ± 1.2% | 33.6 ± 2.4% |
| 1 carbon | 4.3 ± 0.7% | 3.3 ± 0.3% |
| 2 carbons | 45.4 ± 0.6% | 59.3 ± 2.2% |
| 3 carbons | 0.9 ± 0.5% | 1.0 ± 0.5% |
| 4 carbons | 5.2 ± 0.3% | 2.3 ± 0.1% |
| 5 carbons | 0.0 ± 0.1% | 0.0 ± 0.0% |
| 6 carbons | 2.7 ± 0.1% | 0.3 ± 0.2% |
| 7 carbons | 0.0 ± 0.0% | 0.0 ± 0.0% |
| 8 carbons | 0.5 ± 0.2% | 0.1 ± 0.1% |
| 9 carbons | 0.0 ± 0.0% | 0.0 ± 0.0% |
| 10 carbons | 0.0 ± 0.0% | 0.1 ± 0.1% |

